# The canonical HPA axis contributes to locomotion during photoadaptation but is not required

**DOI:** 10.1101/2022.05.14.491921

**Authors:** Han B. Lee, Viet Ha Dang Thi, Grace E. Boyum, Rodsy Modhurima, Emma M. Hall, Izzabella K. Green, Elizabeth Cervantes, Soaleha Shams, Karl J. Clark

## Abstract

The hypothalamic-pituitary-adrenal (HPA) axis and its effector molecules—glucocorticoids—modulate diverse aspects of physiology in vertebrates ranging from metabolism to immune function, behavior, and circadian cycle. While the glucocorticoid receptor (*nr3c1*) is known to be involved in light adaptation (photoadaptation) of the retinal cells, the role of *nr3c1* and other HPA axis signaling molecules in the behavioral phenotypes observed during photoadaptation have not been delineated. Therefore, we investigated locomotor adaptation to various light/dark durations using larval zebrafish that carry a mutated allele in key HPA axis receptors. First, we established baseline locomotion and compared WT and *nr3c1* mutant larvae in constantly lit and dark conditions (for 12-hrs). WT larval zebrafish showed highest locomotor activity in the middle of the day and low activity during the early and later parts of the day, which was generally higher in the light. The baseline locomotion was depressed in *nr3c1* mutants throughout the day in both environments with a more significant change noted in the light. Next, groups of larvae mutant in *nr3c1, nr3c2* (mineralocorticoid receptor), or *mc2r* (melanocortin receptor type 2; adrenocorticotropic hormone (ACTH) receptor) along with their wildtype (WT) siblings were acclimated in the dark and underwent four cycles of dark-light illumination changes with different durations of illumination: 7.5, 6, 4, or 2 min. The *nr3c1* and *mc2r* mutant fish showed significantly decreased locomotion during the dark phase when the illumination was provided for 4 or 2 min. However, with 4 min or longer illumination, if locomotor deficits were observed in mutant larvae, they demonstrated a “catch-up” phenotype with increasing locomotion during the later dark phases of the assay and ultimately reaching the swimming distances indistinguishable from their WT siblings in the assays with 7.5-min illumination. Finally, we looked at effects of light intensity, beginning with our short light exposure. A lower intensity of light failed to elicit any response even from WT fish after 1-min illumination. However, this dim light still evoked a robust response from *nr3c1* mutants and their WT siblings after 7.5-min illumination. The *nr3c2* mutant larvae consistently showed locomotor response similar to their WT siblings regardless of the length of the light phase. Thus, activation of the canonical HPA axis (i.e. *nr3c1, mc2r*) was necessary to induce a rapid locomotor adaptation following after light to dark transition with shorter light exposure times (~≤4 min), but it was not needed for locomotor responses observed after longer exposure to light (~> 4 min). Moreover, the locomotor response after transitioning to dark from a short (1 min) exposure was dependent on the intensity of light, whereas the longer exposure was not. Together, these findings suggest that locomotion after longer exposure to light is not dependent on the HPA axis, suggesting that either parallel independent pathway(s) are responsible for the locomotor response in these cases or that the HPA axis facilitates a primary pathway to increase sensitivity to light exposure.

## Introduction

The hypothalamic-pituitary-adrenal (HPA) axis in vertebrates mediates diverse adaptive processes, collectively called the stress response (SR), that involve metabolism, immune function, behavior, and circadian rhythm.^1–3^ Through the processes in SR, organisms adapt to changes in their body and in the environment to maintain homeostasis. In vertebrates, SR is initiated by the secretion of the corticotropin releasing hormone (CRH) from the paraventricular nucleus in the hypothalamus, followed by the secretion of adrenocorticotropic hormone (ACTH) from the anterior pituitary, resulting in the release of glucocorticoids from the adrenal gland in a cascade.^4^ Glucocorticoids—cortisol in humans and zebrafish and corticosterone in rodents and birds—effectuate SR by binding to corticosteroid receptors (CRs) in cells and terminate SR by providing negative feedback to the hypothalamus and pituitary. The transmembrane ACTH receptor or melanocortin receptor type 2 is encoded by the gene *mc2r*, expressed in the adrenal gland, and initiates the synthesis and secretion of glucocorticoids.^5^ Glucocorticoids signal through at least two key nuclear hormone receptors^6,7^ that differ in their cognate ligands, expression, and affinity to cortisol. The mineralocorticoid receptor (MR), which is encoded by the gene nuclear receptor subfamily 3 group C member 2 (*nr3c2*) shows more tissue-restricted expression including the brain and kidney in mammals, while the glucocorticoid receptor (GR), encoded by *nr3c1* shows more ubiquitous expression in most cells in vertebrates^8–12^. MR has a higher affinity for glucocorticoids like cortisol, is mostly saturated at baseline, and is therefore responsible for setting tonal responses, while GR is thought to be primarily responsive when cortisol spikes with circadian rhythm or acute stress.^11,13,14^

In zebrafish, the hypothalamic-pituitary-interrenal (HPI) axis is functionally homologous to the mammalian HPA axis with the only notable difference being the loose organization of interrenal cells in the kidney versus cell layers found in the mammalian adrenal gland.^15,16^ Zebrafish are quite suitable for behavioral research including light and HPA axis, in part because they are a diurnal species with the same circadian pattern as humans and because of viability of *nr3c1* homozygous mutants^17^. Larval zebrafish develop rapidly establishing most organ systems by 5 days post fertilization (dpf) including the visual system.^18,19^ The genes relevant to HPI axis functions are expressed around the time of hatching (2.5 dpf), and the HPI axis responds to exogenous stressors from 4 dpf onward.^16,19–22^ Larval zebrafish exhibit a repertoire of phototaxis and prototypical swim behavior.^23,24^ Larvae (5 dpf) prefer mildly lit environments, swim toward a lit spot with low intensities, and search for light when placed in darkness (positive phototaxis).^24–26^ When suddenly transitioned from dark-to-light, larvae show a temporary decrease in locomotion and gradually increase locomotion back to baseline levels for lit environments in about 5 minutes. Conversely, a sudden transition from light-to-dark evokes an immediate increase in locomotion that lasts about 10 minutes, after which larvae go back to lower levels of basal locomotion for darkness.^27^

Responsiveness to changing light and photoadaptation are vital for vertebrate homeostasis and the hypothalamus is involved in many aspects of photoadaptation. In vertebrates, light signals are transmitted to the suprachiasmatic nucleus (SCN) of the hypothalamus via the retinohypothalamic tract and the direct innervation of the retinal ganglion cells to the SCN is the basis for the circadian rhythm.^28^ Feeding and metabolism are attuned to the circadian cycle and glucocorticoids synchronize the periphery with the central clock.^29–33^ The glucocorticoid receptor, encoded by *nr3c1*, regulates gene expression in the retina, affecting visual acuity and adaptation of retinal cells during dark to light transitions.^34,35^ However, the extent to which the HPA/I axis modulates adaptive locomotor behavior of animals during light changes remains largely unknown.

Exploiting larval photo responses and a commonly used behavioral assay paradigm, we previously studied the role of HPI axis in SR as larvae (5 dpf) adjust their locomotion in reaction to brief light stimulus. We found that *mc2r* and *nr3c1* homozygous mutant larvae showed significantly decreased locomotion after a brief 1-minute white light when compared to wild-type siblings. In contrast, *nr3c2* homozygous mutants had locomotor response equivalent to wildtype (WT) siblings.^15^ Thus, we concluded that locomotor response to sudden light changes depends on HPI axis activation (*mc2r* and *nr3c1*) and speculated that *mc2r* and *nr3c1*, but not *nr3c2*, would also play a role in broader photoadaptive behaviors. Interestingly, a contemporary report indicated that *nr3c2* was essential for stress axis regulation in larvae, including an altered behavioral response to repeated light changes to longer light stimuli (7.5 min) while *nr3c1* mutants had no effect (Faught and Vijayan, 2018). Thus, the relationship between various key components of HPI axis activation and the elicited locomotion during illumination changes required further investigation.

To better understand the roles *nr3c1, nr3c2*, and *mc2r* play in SR following light changes, we investigated locomotor response (swim distance) to various stimuli of changing light, focusing on larvae homozygous mutant in *mc2r, nr3c1*, or *nr3c2*. We established basal locomotion in WT and *nr3c1* mutant larvae in continually lit and dark conditions for 12-hrs. Without the trigger of sudden light transition, the baseline activity of *nr3c1* mutant larvae was significantly lower compared to their WT siblings in both environments with a greater change observed during the lit condition. Next, based on previous reports and our pilot work, various durations of light (7.5, 6, 4, or 2 minutes) were employed to study locomotion in dark-acclimated fish as they were subjected to repeat dark-light assays. Evaluation of longer light exposure demonstrated that homozygous mutant zebrafish larvae in *mc2r, nr3c1*, or *nr3c2* all swam distances comparable to their WT siblings during the dark phase after 7.5-minute light exposure. The mutant larvae in *mc2r* or *nr3c1*, but not *nr3c2*, showed a decreasing trend of locomotion during darkness, reaching significantly decreased levels of locomotor response in all dark phases in the 2-minute white light regimen. As illumination times increased, phenotypes that were most prominent in the first dark phase with the locomotor response “catching up” to wild type siblings in subsequent light to dark transitions slowly went away with all but the first dark phase in *mc2r* appearing like WT siblings. The locomotor response following the light-to-dark transition that occurred after 7.5-minute illumination was dependent on both the sudden light-to-dark transition and the sufficient length of illumination during the previous step (e.g. 7.5 minute). With these longer exposure times, locomotor response was not dependent on intact HPI axis signaling. Finally, we examined the impact of light intensity on locomotor response. With a one-minute light exposure, a lower intensity of light failed to evoke any significant swimming behavior even from WT larvae while the same low intensity light still elicited robust locomotor response after 7.5-minute illumination in *mc2r* mutants and *nr3c1* mutants along with their WT siblings.

Therefore, we report that the canonical HPI axis (*mc2r, nr3c1*) was necessary for photoadaptive increased swim activity until light was provided for longer durations. Our findings point to two non-mutually exclusive hypotheses that there are one or more parallel pathways achieving photoadaptive behavior in addition to the canonical HPI axis, or HPI axis activity plays a facilitative or permissive role potentiating other indispensable pathways for such adaption.

## Materials and Methods

### Materials and equipment

We listed the materials and equipment used in this study in the supplementary table 1.

### Zebrafish husbandry

Wild-type (WT) zebrafish (*Danio rerio*) originally purchased from Segrest Farm in Florida were outbred to keep a genetically diverse healthy stock. Fish were handled and cared for following standard practices^36^ and guidelines from the Institutional Animal care and Use Committee (IACUC) in the Mayo Clinic (A345-13-R16, A8815-15). Adult fish were kept in a 9 L (25-30 fish) or 3 L (10-15) housing tank at 28.5°C with a light:dark (14:10) cycle. All experiments in this study were conducted before the zebrafish sex can be determined at about 15 dpf,^18,37–39^ and thus sex determination was not made.

### Mutant zebrafish lines

The same WT and mutant zebrafish lines that we previously reported were used.^15^ All fish were maintained through outbreeding. There were three *mc2r* mutant lines that each carried a frameshift mutation in exon 1 (two 4- and one 5-base pair deletions; *mn^57^, mn^58^*, and *mn^59^*, respectively). The annotation on the gene *mc2r* in the National Center for Biotechnology Information (NCBI) has changed from having 2 exons to having 1 exon since our previous report. Four *nr3c1* frameshift mutants were used, each of which carried a 7- or a 17-bp deletion in exon 2 (*mn^61^, mn^62^*) or a 4- or a 5-bp deletion in exon 5 (*mn^63^*, *mn^65^*). A nr3c2 frameshift mutant was used that carried a 55-bp deletion in exon 2 (*mn^67^*). For detailed information on the mutant lines, refer to our previous paper ^15^.

### Custom light boxes

Light boxes were custom produced by the Mayo Clinic Division of Engineering. The light box has a control panel with two knobs that enable light intensity adjustment—both white light and infrared (IR) light have a low, medium, and high intensity. The intensities for white light were ~8,000, ~4,000, and ~300 lx for the high, medium, and low setting, respectively. Infrared illumination does not produce any light power meter reading (0 lx). The high intensity was used for both white and IR light. For the dim light assays, the low intensity was used. For detailed information on the custom light boxes (e.g. dimensions, emission spectral graphs of light), refer to our previous paper ^15^.

### Behavioral assays preparation

An adult mating pair were placed in a mating tank separated by a divider (−1 dpf). On the following day, the divider was pulled in the morning and embryos were obtained via natural spawning (0 dpf). Unfertilized, dead, or morphologically defective embryos were cleaned up on 0, 1, and 3 dpf. On 3 dpf, a single larva was placed in each well of a 48-well plate. On 5 dpf, behavioral assays were performed in a custom-built chamber (Fig. 1). From day 0-5, plates were stored in an incubator with a light:dark cycle (14:10 hrs) at 28.5°C.

**Figure 1.**
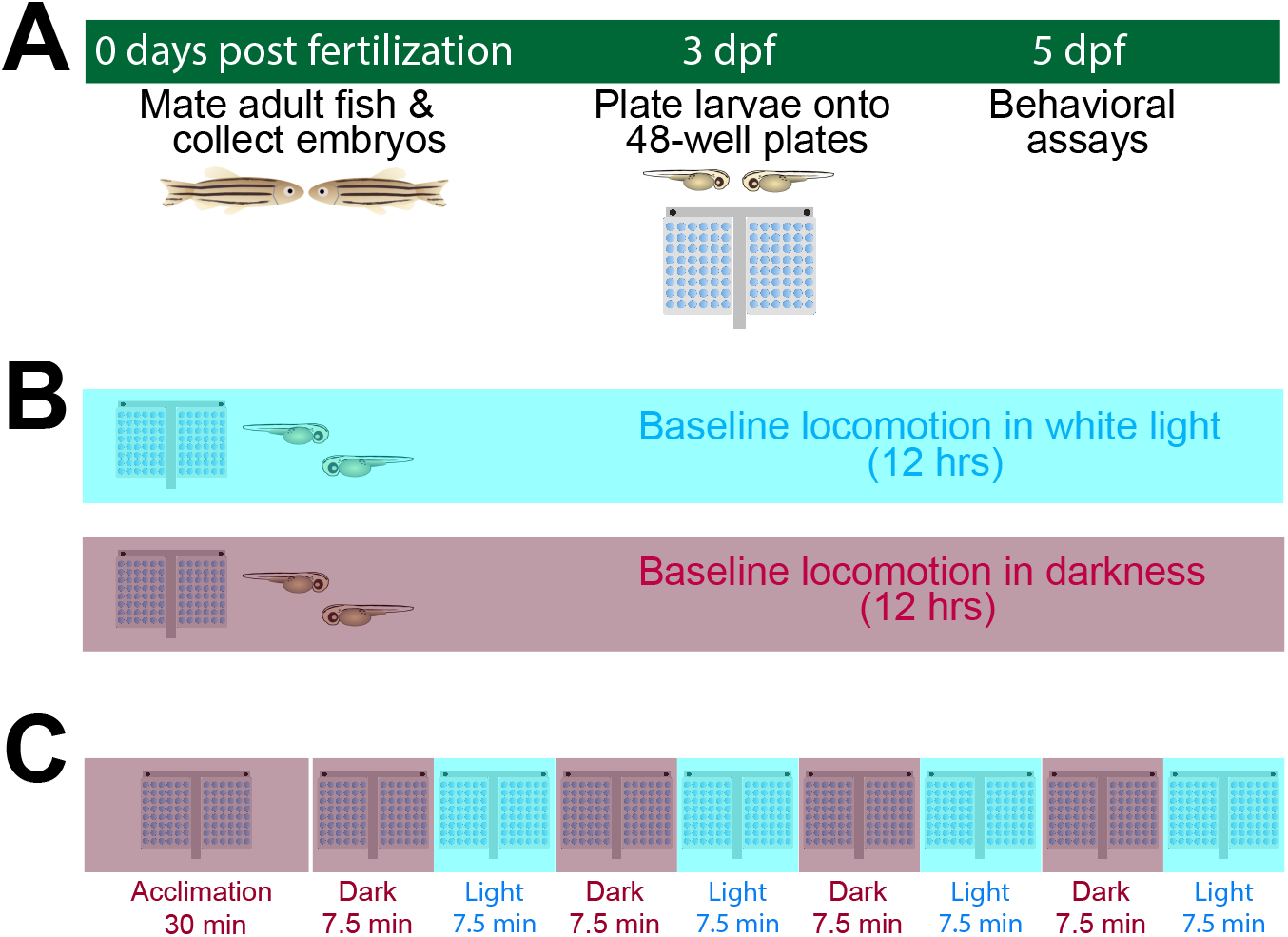
Experimental processes and behavioral assay paradigm. (A) Experimental preparation. Embryos were obtained via natural breeding. Fertilized embryos were collected on 0 dpf (days post-fertilization) and kept in an incubator at 28.5°C. Dead and poorly developing embryos were cleaned on 0 and 1 dpf. Individual larvae were placed (one per well) in a 48-well plate on 3 dpf. (B) Basal locomotor assay paradigm. Fish were acclimated to dark or light for 30 min and then video recorded for 12-hrs on 4, 5, 6 or 7 dpf. Locomotor activity in the dark was recorded in Infrared light (IR, 0 lx) and activity in the lit environment was recorded in white light (~ 8000 lx). (C) 7.5-min dark-light repeat assay paradigm. Behavioral assays were performed on pfd 5. Fish plated on 3 dpf were acclimated to dark and recorded for 60 min with alternating dark-light phases. Dark condition was recorded in IR (0 lx) while lit condition were recorded in diffused white light (~ 8000 lx).

### Basal locomotor activity assays

The basal locomotor assays were performed without light changes. After fish were prepared for assays, they were acclimated in dark or light for 30 minute before videorecording started. Videorecording started around 9 am and the recording continued for 13 hours without any other changes from the light condition that the fish were acclimated (Fig. S1). Larvae were tested at 4, 5, and 6 dpf.

### Dark-light repeat assays

The dark-light repeat assays were performed, using the light illumination changes as an exogenous stressor. The dark period is recorded in infrared (IR). Although zebrafish are thought to be unable to detect infrared, negative phototaxis was also reported.^40^ During the light period, diffused white light illumination was provided from the bottom. The intensity (~8,000 lx) was similar to regular indoor lighting levels (Fig. 1). The locomotor response of the fish was videorecorded.

The larval zebrafish on 5 dpf were acclimated in a dark environment for 30 minutes and underwent the dark (7.5 min) and light (7.5 min) periods four times. In the assays with shorter durations of illumination, 2, 4, or 6-min illumination was used while keeping the length of the dark phase constant at 7.5 minutes. Regimen: 30-min dark acclimation – 4x [7.5-min dark – 7.5-min light] (Fig. 1).

### Dim-light assays

In dim light assays, the same protocol was followed except that the lowest of three intensity settings was used for white light (~300 lx, compared to ~8,000 lx for most assays).^15^ Dim-light assays were performed with 1-min and repeat 7.5-min light exposure.

### Statistical analyses

Reported data are means ±95% confidence interval (CI) unless otherwise stated. Statistical analysis was performed using R language.^41^ For multiple statistical comparisons, multivariate two-way mixed analysis of variance (multivariate mixed ANOVA) was used. A between-subjects factor was the genotype of the fish (i.e. wildtype (WT), heterozygous, or homozygous) and a within-subjects factor was the changing illumination at different time points (i.e. 8 phases; dark—light—dark–light—dark—light—dark—light). When Mauchly’s sphericity assumption was violated, Greenhouse-Geisser correction was applied for an appropriate correction. After the correction, the adjusted *F* value, *p* value, and degrees of freedom were reported. ANOVA analyses were followed by post-hoc analysis (Tukey’s honest significant difference test) to identify the components where the significance arose. For two group comparisons, the Student’s *t* test was used. To improve the readability of the paper, the Result section was written in colloquial language while reporting the key findings. Rigorous statistical analyses and precise numbers are fully reported in the **Supplemental Result 1**.

## Results

### Loss of *nr3c1* larvae exhibited decreased basal locomotor activity in lit and dark environments

While we noticed that *mc2r* and *nr3c1* homozygous mutants tended to move less even during the light phase in our previous investigation, the reduction in locomotion did not reach statistical significance. We were curious whether some of the apparent decrease of locomotion in *mc2r* and *nr3c1* mutants arose from the 1-min light exposure or was reflective of lower baseline activity levels.

To investigate the questions, we quantified the basal levels of locomotor activity of *nr3c1* WT, heterozygous, and homozygous siblings, as well as the age-matched larvae from the general WT stock in the lab, in the lit or dark conditions without any other environmental changes. Larvae from the general WT stock showed a circadian variation in locomotor activity with the highest activity in the middle of the day (Fig. 2A). Larval movement increased each day, with locomotor distance being the greatest at 6-dpf (Fig. 2B). Homozygous mutants in *nr3c1* exon 5 showed decreased basal locomotor activity throughout the day in both lit and dark conditions (Fig. 1B). The reduction in basal locomotion reached significance in some segments of the day (e.g. middle of the day) and was more apparent when lit. The results showed that the decreased baseline locomotion in nr3c1 is a phenotype dependent on the genotype.

**Figure 2.**
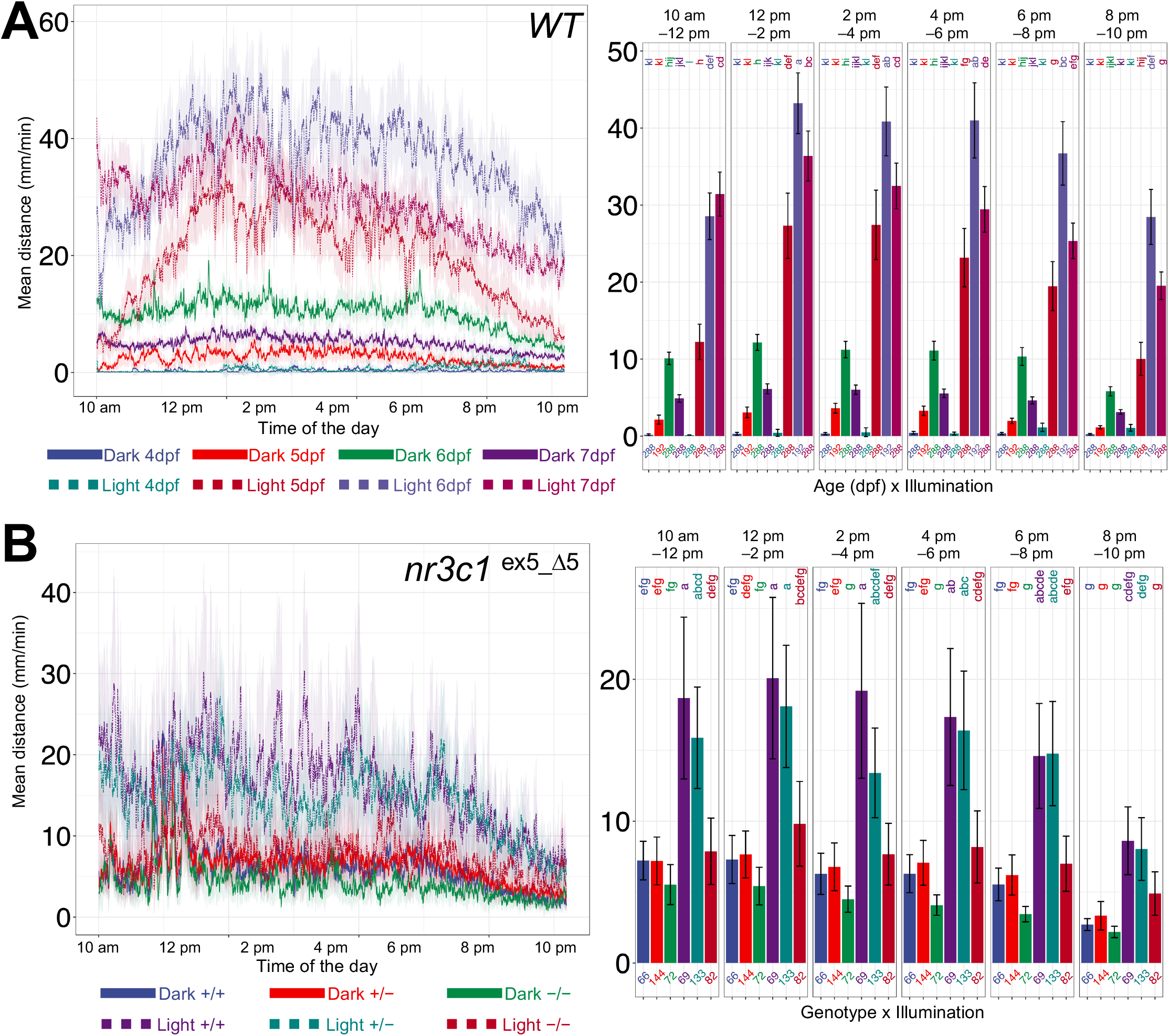
Loss of *nr3c1* larvae exhibited decreased basal locomotor activity in lit and dark environments. (A) Basal locomotor activity in WT larvae over 12-hrs. Zebrafish larvae (4, 5, 6, and 7 dpf) show increased locomotor activity in a lit environment, compared to their counterparts in the dark. In the dark, larvae (4 and 5 dpf) show minimal locomotor activity while 6-dpf larvae swim marginally but significantly more. 6-dpf larvae swim the most distance in the lit condition. The larval zebrafish show increased locomotor activity in the middle of the day and decreased locomotion during the early and later of the day. Line graphs show the mean distance moved (averaged over one-min). Bar graphs show the mean distance larvae moved over 2-hr periods (mean ± 95%CI) with different letters indicating a significant difference between groups and/or different 2-hr periods (Tukey’s honest significant difference test, p < 0.05). Sample size for each condition is shown at the base of each bar. (B) Basal locomotor activity in larvae homozygous mutants in *nr3c1* exon 5 over 12-hrs. Homozygous *nr3c1* mutant (6 dpf) show significantly decreased locomotor activity during the time when WT siblings are most active (middle of the day) in the lit environment. Homozygous larvae move consistently less than their WT and heterozygous siblings, but did not reach statistical significance. Larvae were derived from crosses of *nr3c1^+/−^* fish.

### Mutant *mc2r* and *nr3c1* larvae showed locomotion equivalent to their WT siblings in 7.5-min repeat assays

We previously reported that locomotor response to sudden light changes depends on HPA axis activation in the 1-min light assay (dark acclimation – 1-min white light – 30-min dark).^15^ Considering the relation between the glucocorticoid receptor and light adaptation,^34,35^ we asked whether increasing the duration of the light phase alters the phenotype and adapted a locomotor assay paradigm commonly used in the zebrafish community to test the HPI receptor mutants.^40,42,43^ After being acclimated in the dark for 30 min, larval zebrafish (5 dpf) underwent 7.5-min dark/light repeats four times (Fig. 1B). Unlike after a one-min exposure to light, the *mc2r* and *nr3c1* homozygous mutants swam distances indistinguishable from their wildtype (WT) siblings during the dark phase (Fig. 3). Mutant *mc2r* larvae showed reduced locomotor activity during the dark phase following the first light-to-dark transition but increased activity in the later cycles, showing a “catch-up” phenotype (Fig. 3A). Since no other conditions except the duration of illumination were modified between the 1-min and 7.5-min assays, we asked at what duration of light exposure the locomotor deficit in *mc2r* and *nr3c1* mutants became apparent.

**Figure 3.**
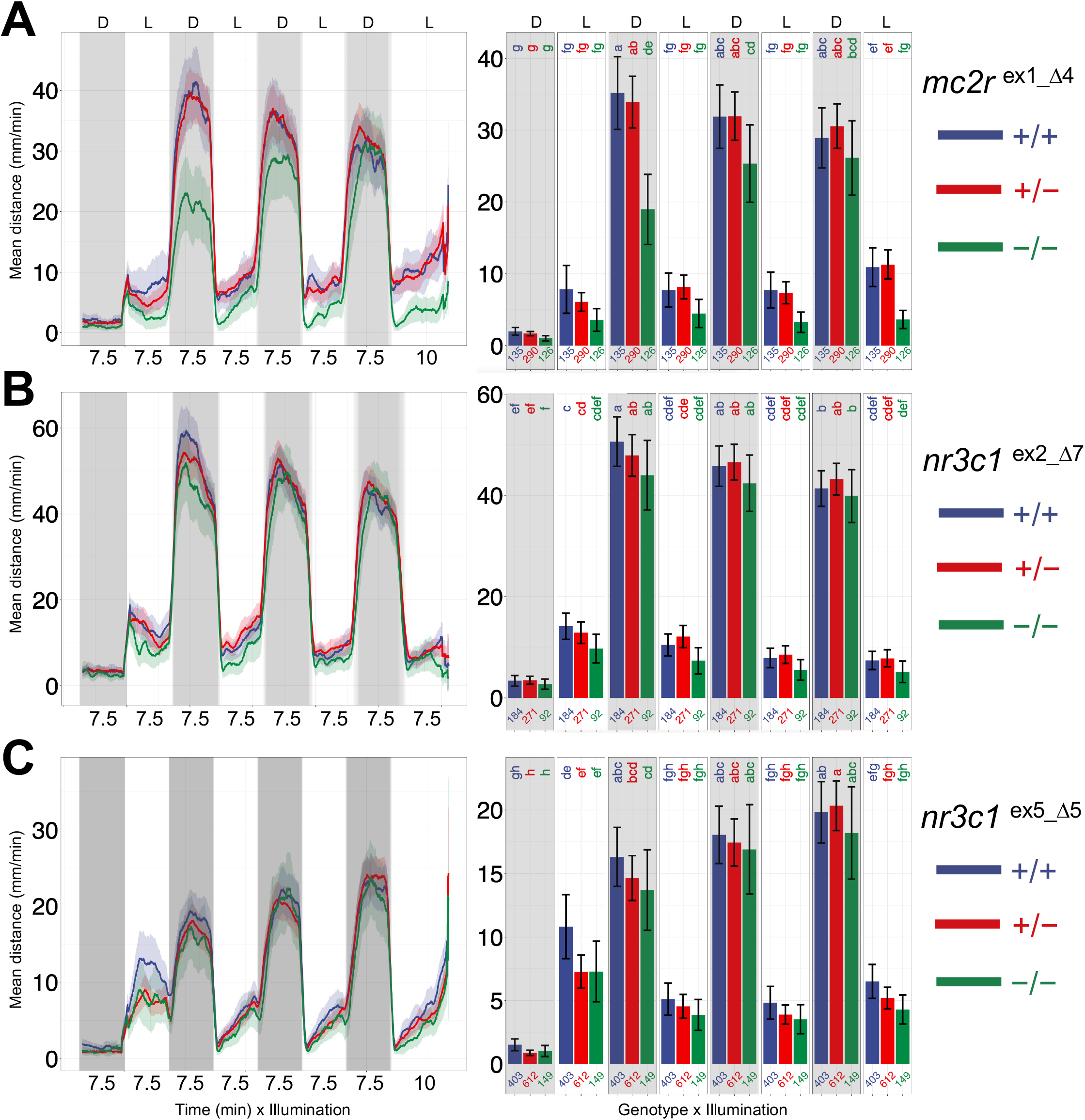
Mutant *mc2r* and *nr3c1* larvae showed locomotion indistinguishable from their WT siblings in 7.5-min repeat assays. (A) Homozygous mutants in *mc2r* increase locomotor response during the dark phase in the later cycles, reaching equivalent levels of swim distance as that of WT siblings. Larvae were derived from crosses of *mc2r*+/− fish (WT^+/+^, heterozygous^+/-^ or homozygous^−/−^). Line graphs show the mean distance larvae (5 dpf) moved. Locomotor activity at each point is the mean distance fish moved during the preceding 60 seconds (mean ± 95%CI [shading]). Bar graphs show the mean distance larvae moved over the time course (mm/min; mean ± 95%CI). Sample size for each condition is shown at the base of each bar. (B, C) Homozygous mutants in *nr3c1* exon 2 and exon 5 show locomotor activity comparable to WT siblings in the dark phase, respectively.

### Mutant *mc2r* larvae exhibited decreased locomotion in 4- or 2-min light illumination

We altered the paradigm by shortening only the duration of white light exposure. When the duration of light was 4 or 2 min, mc2r homozygous mutants showed significantly decreased locomotor activity compared to the WT siblings (Fig. 4).

**Figure 4.**
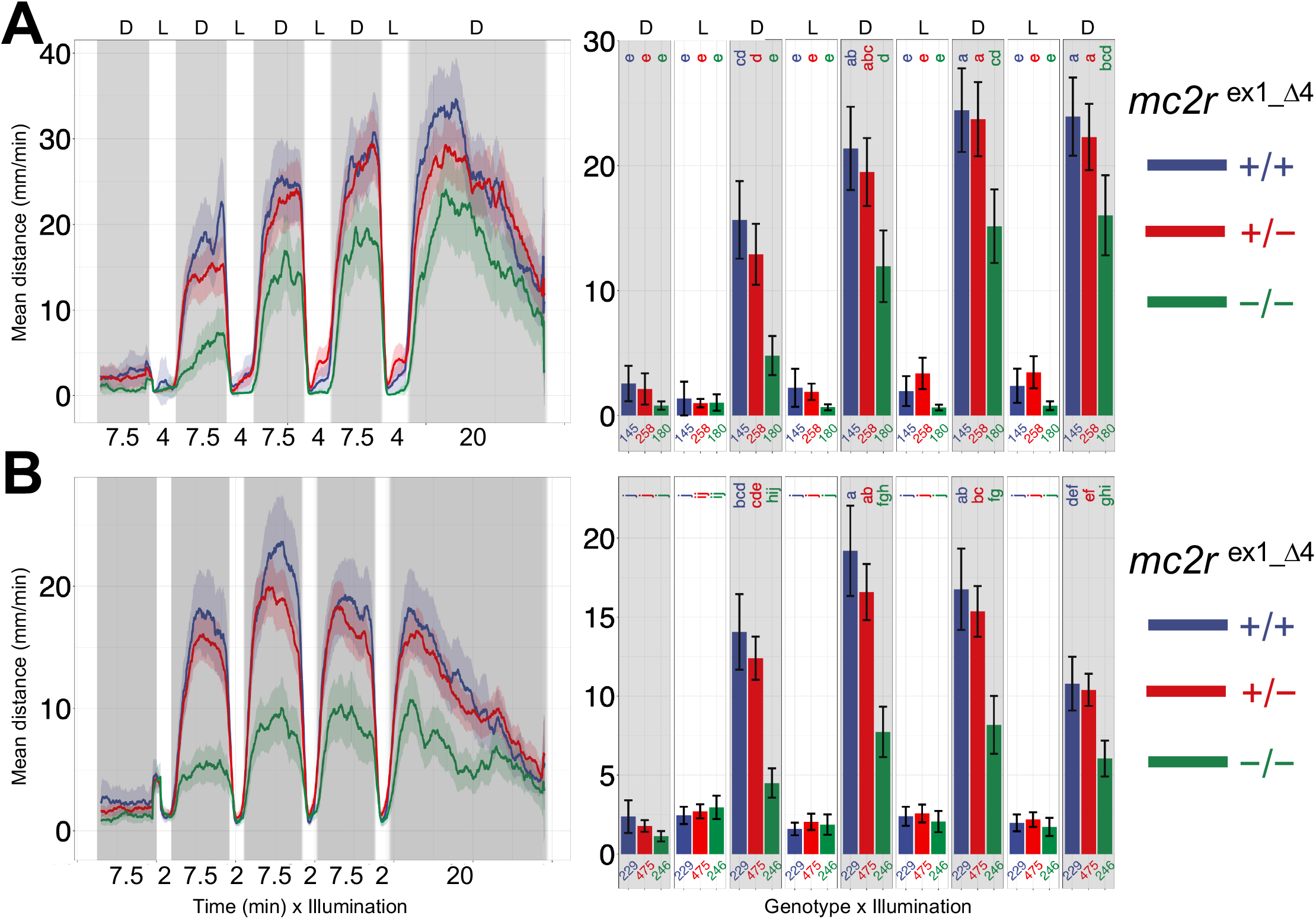
Mutant *mc2r* larvae exhibited decreased locomotion in 4- or 2-min light illumination. (A) *mc2r* homozygous mutants show decreased locomotor activity during the dark phase while showing gradual increase toward the later cycles in the dark (7.5 min)—light (4 min) repeat assay. (B) *mc2r* homozygous mutants display decreased locomotor activity during the dark phase in the dark (7.5-min)—light (2-min) repeat assay.

### Mutant *nr3c1* larvae showed decreasing trends in locomotion when subjected to 6-, 4-, or 2-min of light

When the duration of light was 6 min, homozygous mutants in *nr3c1* exon 2 and exon 5 showed “catch-up” phenotypes, with an initial deficit in locomotor response relative to WT siblings that approached and caught WT locomotor levels during the dark phase in later cycle. However, different from *mc2r* mutants, the depression of the locomotor response in *nr3c1* mutants during the earlier cycles (1^st^ and 2^nd^) of the dark phase was not significant albeit a clear decrease (Fig. 5A, 6A). The 4-min and 2-min illumination led to significantly decreased locomotor activity during the dark phase in *nr3c1* homozygous mutants (Fig. 5B, 6B, and 6C). Thus, the levels of locomotor activity varied based on the duration of the light.

**Figure 5.**
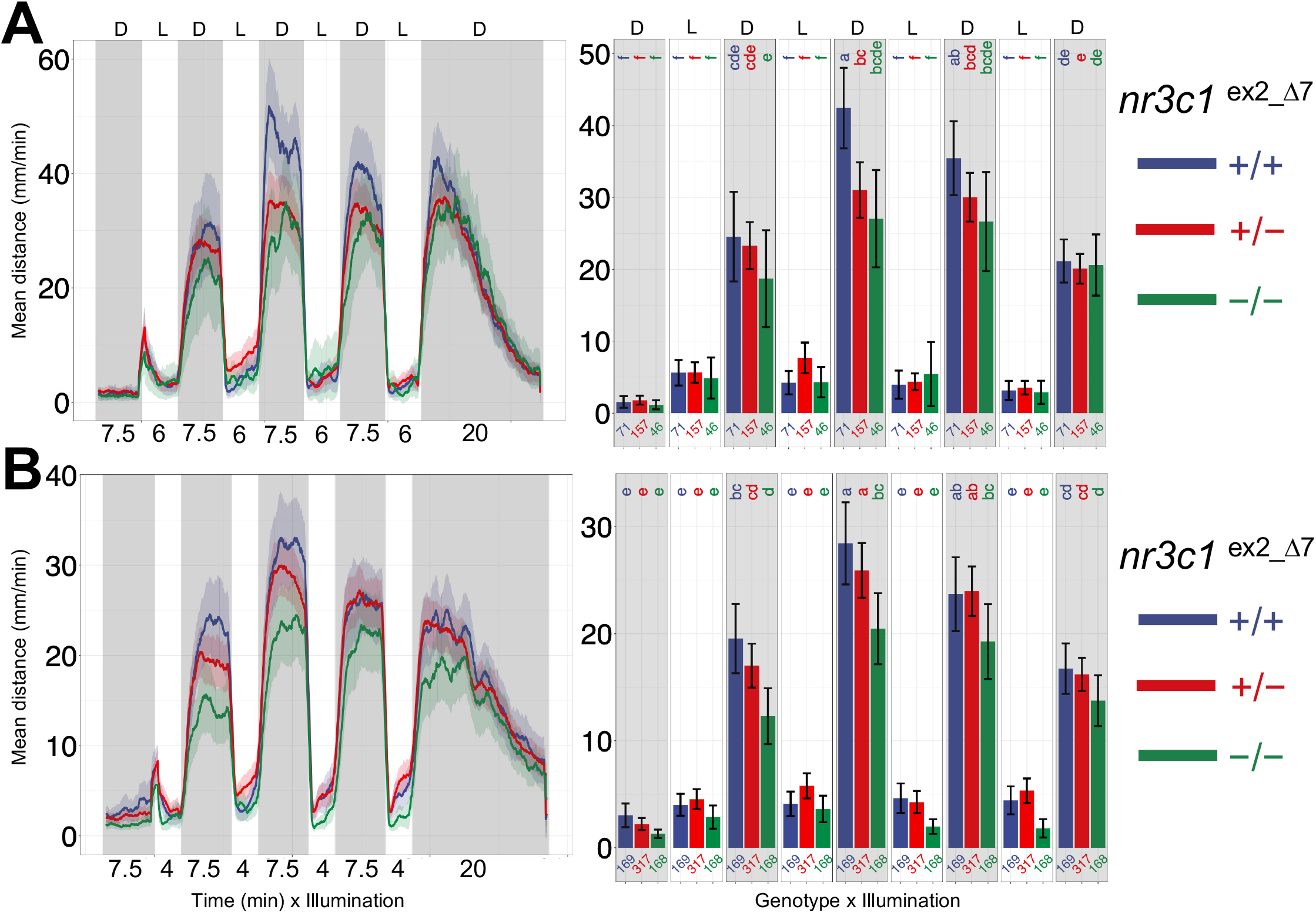
Mutant *nr3c1* (exon 2) larvae showed decreasing trends in locomotion when subjected to 6-, or 4-min of light. (A) Homozygous mutants in *nr3c1* exon 2 show locomotor activity equivalent to WT siblings during the dark phase in the dark (7.5 min)—light (6 min) repeat assay, except the third dark phase. (B) Homozygous mutants in *nr3c1* exon 2 exhibit significantly decreased locomotor activity vs. WT siblings during the first and second dark phases in the dark (7.5 min)—light (4 min) repeat assay.

**Figure 6.**
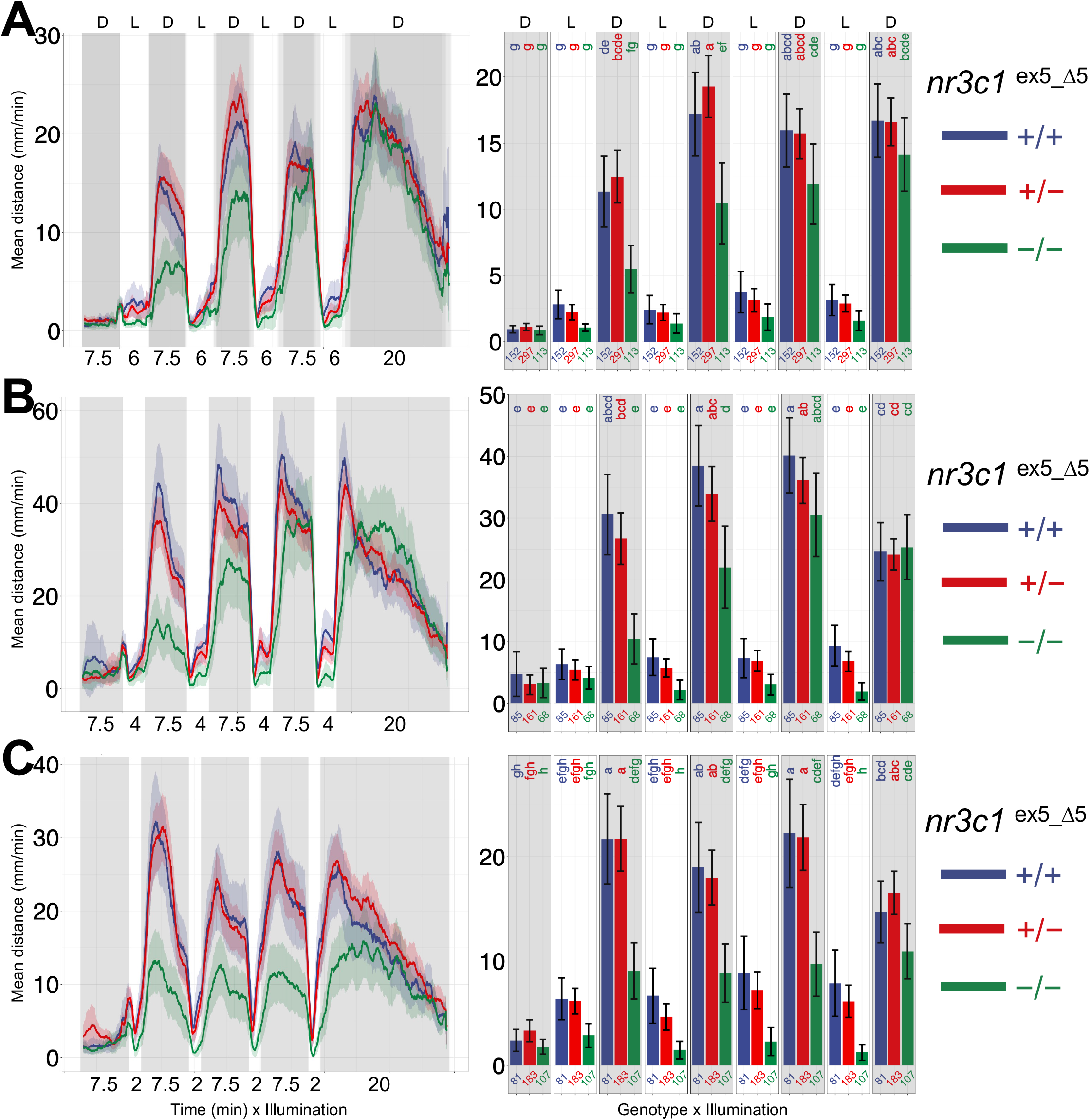
Mutant *nr3c1* (exon 5) larvae displayed decreasing trends in locomotion when subjected to 6-, 4-, or 2-min of light. (A, B, and C) Homozygous mutants in *nr3c1* exon 5 show significantly decreased locomotor activity compared to WT siblings in at least several dark phases in the dark (7.5 min)—light (6, 4, and 2 min, respectively) repeat assay.

After confirming the light dose-dependency of the locomotor phenotype, we asked two questions. One, whether the other key corticosteroid receptor—*nr3c2*—is also involved in this phenotype since the 1-min paradigm elicited no phenotypes in *nr3c2* homozygous mutants. Two, whether *mc2r* and *nr3c1* mutants have basal locomotor activity equivalent to WT siblings in extended lit or dark environments.

### Mutant *nr3c2* larvae showed locomotion similar to their WT siblings in 7.5-, 4- or 2-min illumination

When tested with 7.5-, 4- or 2-min light illumination, homozygous mutants in *nr3c2* exon 2 showed locomotor activity similar to their WT siblings (Fig. 7). 7.5-, 4-, 2-, and 1-min light did not evoke any distinct phenotypes in *nr3c2* mutants. Although there are marginally decreasing trends in locomotion after 2-min illumination (Fig. 7C), *nr3c2* did not appear to significantly contribute to the light adaptation process tested in the dark-light assay paradigm.

**Figure 7.**
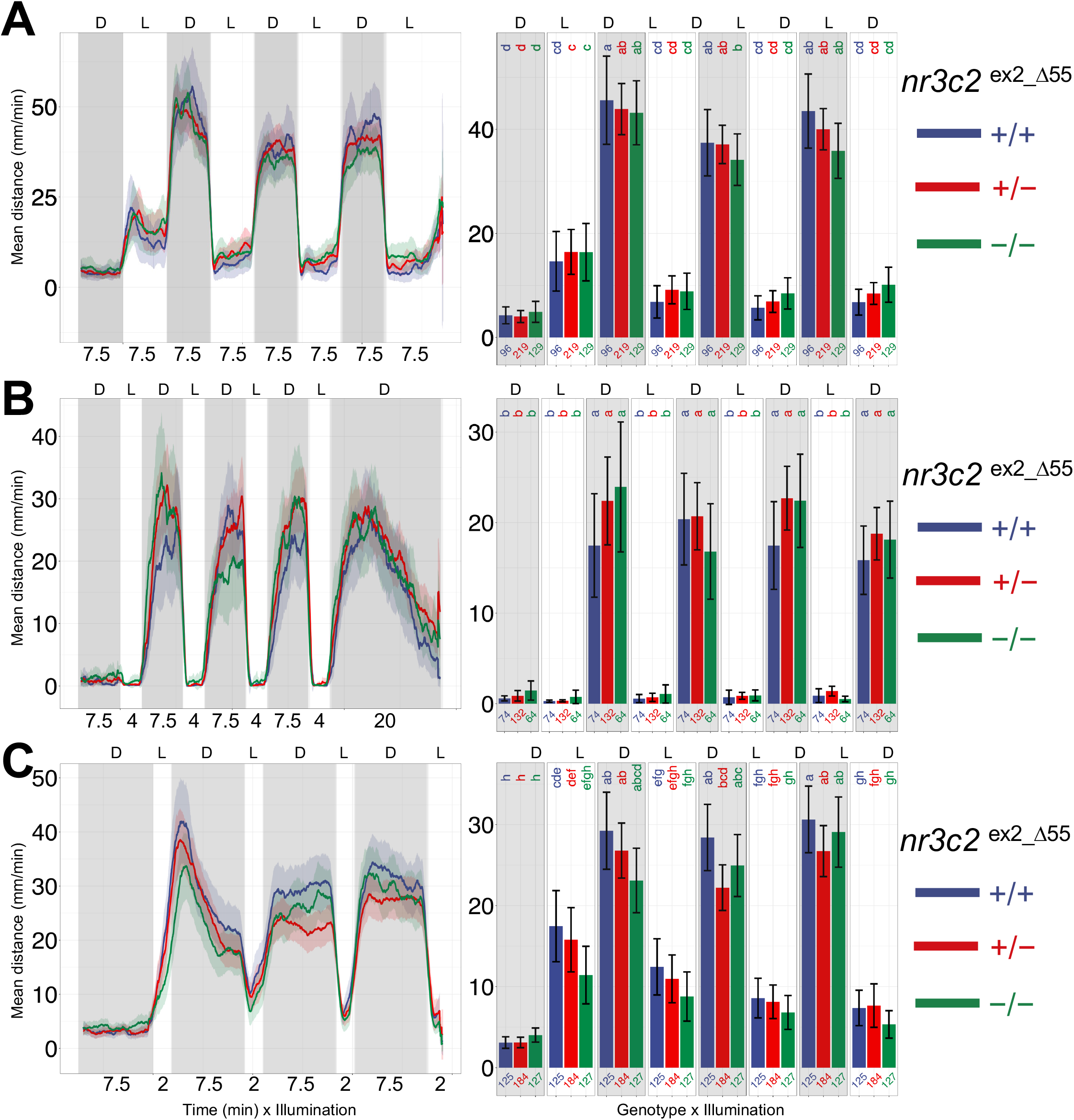
Mutant *nr3c2* larvae showed locomotion similar to their WT siblings in 7.5-, 4- or 2-min illumination. (A, B, and C) Homozygous mutants in *nr3c2* exon 2 displayed locomotor activity equivalent to WT siblings during the dark phase in the dark (7.5 min)—light (7.5, 4 and 2 min, respectively) assay. Larvae were derived from crosses of *nr3c2*^+/−^ fish. Line graphs show the mean distance larvae moved (5 dpf). Locomotor activity at each second is the mean distance fish moved during the preceding 60 seconds (mean ± 95%CI [shading]). Bar graphs show the mean distance larvae moved over the time course (mm/min; mean ± 95%CI).

### Dimmer white light produced similar locomotion in *nr3c1* WT, heterozygous, and homozygous siblings

Since the increased locomotion of *mc2r* and *nr3c1* mutants during the dark phase was triggered by the dark-light transitions and dependent on the duration (quantity) of light illumination, we asked whether the increased locomotor activity also depended on the intensity of light illumination. We tested a low intensity light exposure (~300 lx) to see how it impacted locomotor responses. Following a 1-min dim light exposure, neither mutant nor WT larval fish demonstrated any discernable changes in locomotion (Fig. 8A). When using the dim light for the 7.5-min dark-light repeat assay, *nr3c1* WT, heterozygous, and homozygous siblings all showed robust locomotor activity equivalent to that shown after light illumination of higher intensities during the dark phase (Fig. 8B). Our findings showed that locomotor response after shorter durations (1-min) but not the longer durations (7.5 min) of illumination are sensitive to intensity.

**Figure 8.**
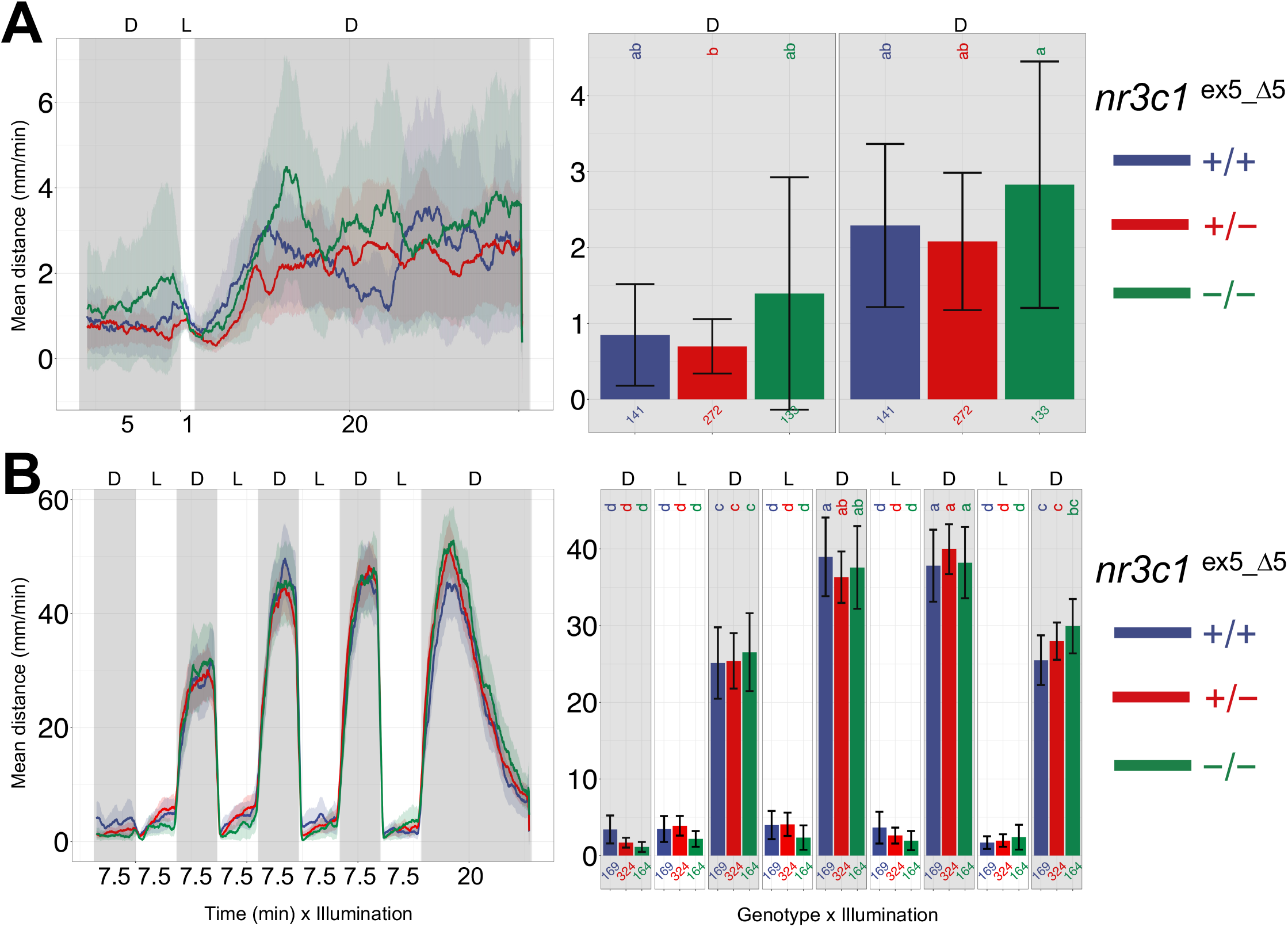
Dimmer white light produced similar locomotion in *nr3c1* WT, heterozygous, and homozygous siblings. (A) Dimmer light (low intensity; ~300 lx) fails to elicit any meaningful pattern of locomotor response from *nr3c1* WT, heterozygous, and homozygous siblings. (B) The same dimmer light (low intensity; ~300 lx) evokes robust and equivalent locomotor activity during the dark phase in *nr3c1* WT, heterozygous, and homozygous siblings.

## Discussion

We had previously demonstrated that exposure to one minute of white light elicited a locomotor response in larval fish. This response was significantly reduced in fish that lacked a functional ACTH receptor (*mc2r*) or GR (*nr3c1*), as compared to wild-type siblings. In contrast, mutations in MR (*nr3c2*) did not result in any loss of locomotor response when compared to their WT siblings. Zebrafish display a wide range of photoadaptation responses. This includes the photomotor response that occurs in 1-2 dpf larvae,^44,45^ phototaxis with a preference towards light,^24^ and the light-to-dark transition.^46^ Even within the light-dark transition space, there are variations in design. For example, we used a short exposure to white light on dark-acclimated larval fish preceding an extended dark phase. Though it is common to use longer phases (7.5-10 minutes) of light and dark and then repeat these 3-5 times.^42,47,48^ In one example of this longer repeating light-dark transition papers, homozygous mutant MR (*nr3c2*) but not GR (*nr3c1*) larval fish had altered locomotor responses when compared to unrelated wild-type fish.^49^ This result contrasted with our one-minute exposure data and suggested there may be differences in photoadaptation signaling depending on assay design-in this case, the length of illumination phase. Therefore, we set out to investigate the influence of HPI axis signaling on photoadaptation, specifically measuring locomotor behavior using different paradigms of light-to-dark transition.

In our previous work with the *mc2r* and *nr3c1* mutant fish subjected to a hyperosmotic challenge, we also noticed a reproducible, but non-significant reduction in baseline locomotion in light-acclimated fish.^15^ These baseline differences were not readily apparent in dark-acclimated fish exposed to white light for one-minute. Here we evaluated extended locomotor recordings in light or dark to fully describe baseline locomotor activity in wild type and mutant fish. All fish demonstrated increased activity in the middle of the recording session (mid-day), but this was more pronounced when fish were recorded in the light. Importantly, mutations in *mc2r* and *nr3c1* decreased baseline locomotor rates, and this was again more pronounced while recorded in light. Therefore, we chose to work primarily with dark-acclimated fish for observing changes due to HPA axis signaling to eliminate background locomotor differences as much as possible.

After acclimating fish in the dark, we implemented a 7.5-minute light exposure followed by 7.5 minutes of dark and repeated this light-dark switch four times. As expected, wild type fish demonstrated a robust increase in locomotion following each light-to-dark transition. In stark contrast to our one-minute exposure assays, there was limited or no locomotor difference between *mc2r* and *nr3c1* mutant fish and their wild-type siblings. Similar to our one-minute white light assays but in contrast to published results,^49^ we saw no locomotor difference between *nr3c2* fish and their wild-type siblings. We suspect this might be due to breeding differences between their study and this one.

It appeared however that the role of the HPA axis in eliciting a locomotor response changes with increasing length of light exposure prior to the dark phase. To discover whether these changes occur at an abrupt time point or gradually change with increasing light exposure, we altered the length of the light illumination phase while keeping the length of the dark phase constant at 7.5 minutes. With decreasing durations of light (7.5, 6, 4, and 2 minutes) locomotor activity became more dependent on HPA axis signaling. As the light duration was lowered, the impact of *mc2r* or *nr3c1* loss was observed in the first light-to-dark transition with subsequent light-to-dark transitions becoming more similar to (catching up) wild-type sibling locomotor levels. As the time was shortened, the impact of the loss of HPA axis signaling became stronger, and when decreased to two-minutes, loss of *mc2r* and *nr3c1* had a strong impact on locomotion in all dark phases. At the population level, decreasing the light illumination phase resulted in a gradual loss of locomotor response in HPA-axis mutants. However, it is possible that the switch was more binary, but the timing of the switch based on sensing light exposure was widespread across the population of fish.

Accordingly, since light exposure may be measured based on the length of exposure (time), intensity of light (lux), or a combination of these two factors, we manipulated the light intensity in both the short light exposure (1-minute) and the repeating dark-light (7.5 minute) paradigm by using a lower intensity light (dim=~300 lx). We observed a dramatic loss of locomotor response in both wild-type and mutant fish exposed to 1-minute of dim light, whereas the dim light had no impact on the repeating dark-light assay, both WT and mutant siblings had robust locomotor responses. Therefore, the short light exposure that was sensitive to loss of HPA axis signaling was also sensitive to light intensity. This observation further helped us solve another mystery. We had purchased commercial behavioral recording stations with an optional bright light plate, and had difficulty reproducing our 1-minute light assays with them. We have since empirically determined that these commercial units required 2.5 to 3 minutes of white light exposure to elicit a locomotor response in wild-type fish. In these assays, locomotion is still impacted by loss of *mc2r* and *nr3c1* (unpublished work not shown).

### Both freezing and increased locomotion may not be related to HPA axis

Larval behavioral studies often rely on locomotor response as a primary quantitative measure in response to various cues. However, locomotion in larval fish, whether decreased or increased, may be due to different parallel signaling pathways that are cued by different stimuli. Although freezing behavior is considered a fear-related response in rodents and adult zebrafish,^50–53^ the nature of freezing behavior in larval zebrafish has not been fully investigated in varying contexts. In our assays, regardless of exposure time, concomitantly with white light being turned on, the zebrafish larvae reduced swimming below baseline, which has been described as a freezing response, and then gradually swam more. The freezing behavior of larvae at the light transition has been associated with fear/anxiety-like behavior and stress response.^49,54,55^ However, our observation indicated that we might need alternative hypotheses to interpret this freezing behavior. Notably, this response still occurred with loss of *mc2r* and *nr3c1*. The sharp reduction of motion at the dark-to-light transition point happened regardless of the genotype of the fish (WT, het, and hom siblings in *mc2r* and *nr3c1;* Fig. 2, 3, 4). Particularly, the reduction was more pronounced in homozygous mutants (*mc2r* and *nr3c1*) compared to their WT siblings. The canonical HPI axis appeared to contribute to the modulation of the drop in motion (zone A in Fig. 9A). Subsequently, the larvae gradually increased locomotor activity levels in lit environments. The pattern of increased locomotion was typical of the baseline locomotion in *nr3c1* WT, het, and hom siblings, in which mutant larvae showed a decreased trend compared to WT (Fig. 2B).

**Figure 9.**
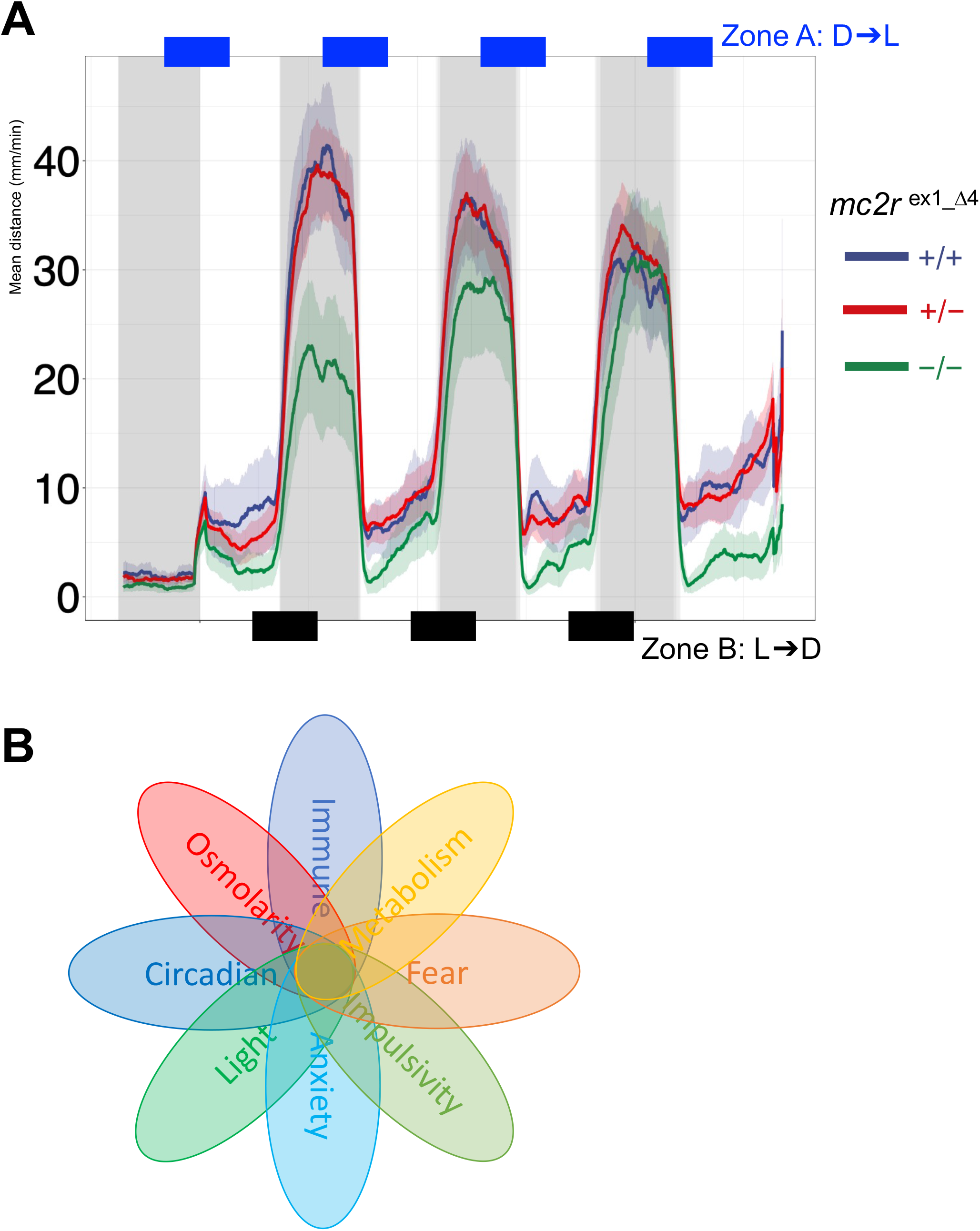
A schematic of light transition. (A) The zone A denotes the dark-to-light transition. Immediately after the illumination change, fish show sharply decreased locomotion. *mc2r* homozygous mutants show a pattern of low activities levels compared to their WT and heterozygous siblings during the light phases, albeit not statistically significant. The zone B marks the light-to-dark transition. Immediately after the transition, fish show sharp increase in locomotion. When light is provided with a sufficient quantity (here, 7.5 min), even HPA axis mutant fish show increased locomotion. (B) A few selected examples of physiological processes that the vertebrate stress response is involved. Each ellipse represents genes and pathways in the process. For example, SR modulates risk-taking behaviors as well as photoadaptive behaviors. The central nervous system-centered processes such as fear-related or anxious behaviors have common components with the metabolic process. Behavioral screening for a specific SR process has a potential to reveal the core program involved in SR.

On the other hand, increased locomotion has also been considered an anxiety-related behavior in rodents and zebrafish depending on the context.^53,55–57^ In our assays, when the light was turned off (zone B in Fig. 9A), larval zebrafish immediately responded with sharply increased locomotion followed by gradual decrease, and returning to baseline in about 15 minutes. We showed that the increase in locomotion at the light-to-dark transition depended on the genotype of fish, and the duration and the intensity of illumination. While WT fish exhibited increased locomotion regardless of the illumination length (Fig. 2, 3, 4),^15^ mutant larvae (*mc2r* and *nr3c1*) displayed increased locomotion only after longer durations of illumination (6- and 7.5-minutes). Since the same stimulus (light-to-dark transition) produced differing locomotor response in shorter or longer durations of light, the illumination transition per se may not be anxiety-inducing. Rather, a physiological process (e.g. accumulation or degradation of cyclic molecules) that occurred during the prior light phase may be responsible for the phenotype with the shorter duration of light. For example, similar delay in the recovery of retinal cell function was found in *nr3c1* mutant zebrafish when light was turned on. These mutant larvae (*gr^s357^*) showed defects in the optokinetic response of the eye due to dysregulation of gene transcription in the retinal cells.^35^

Thus, more work is needed to better understand both freezing and increased locomotion that occur with changing light, in the context of photoadaption, and not only related to fear and anxiety. Since photoadaption is vital for homeostasis, the canonical HPA axis (*mc2r, nr3c1*) is utilized but does not account for the whole locomotor response. Given that the axis is required only after shorter illumination lengths, its activity may be one of multiple pathways that may synergistically facilitate more efficient function with less light. Such modulatory roles of glucocorticoid receptor and other nuclear receptors have been well documented in the investigations on sensitization and potentiation.^58–62^

### Consequences of an HPA axis receptor knockout differ

Seemingly comparable mutations in the HPI axis genes (*mc2r, nr3c1*) may result in different physiological states. Many *nr3c1* mutant animal models have been reported to be hypercortisolemic since the negative feedback loop to the hypothalamus and pituitary was rendered dysfunctional. These mutant animals and patients with mutations in *NR3C1* are sensitized to ACTH, showing the hypercortisolemic state with concurrent paradoxically low ACTH levels.^4,6,14,63–65^ *NR3C1* has at least 5 alternative transcripts and 8 translational isoforms, which is presumed as the basis for the single glucocorticoid ligand (i.e. cortisol) to be highly pleiotropic and have tissue-specific effects^66–68^. It will be important to investigate whether alternative isoforms compensate for the loss of canonical *nr3c1*. In contrast, it is expected that *mc2r* mutant animals were hypocortisolemic and adrenal gland development was inadequate since the canonical ACTH receptor (*mc2r*) is knocked out.^5,69^ Although ACTH-stimulated cortisol production is the primary source of cortisol following an acute stressor, there are potentially other pathways involved in glucocorticoid synthesis. There are several direct innervations of the sympathetic neurons to the adrenal cortical tissue, secondary autonomic innervations from the adrenal medulla, and paracrine effects of the catecholamines secreted from the adjacent chromaffin cells in humans.^70^ In zebrafish, the interrenal area contains both chromaffin cells releasing catecholamines and steroidogenic cells releasing glucocorticoids in close contact.^71,72^ Such anatomical organization would increase the paracrine effects of interrenal cells. Zebrafish Mc4r has been shown to function as an ACTH receptor when melanocortin receptor accessory protein 2 (*mrap2*) was coexpressed ^73^.

In our investigation with various light stimuli, mineralocorticoid receptor (MR, *nr3c2*) did not play a role in photoadaptation. Compared to GR, MR has a ten-fold higher affinity for cortisol and is thought to establish the tonal (basal) function of SR and initiate specific SRs in mammals.^11,13,14^ Teleost fish like zebrafish do not produce aldosterone^74–78^ and osmoregulation is achieved through glucocorticoid receptor (*nr3c1*) signaling rather than *nr3c2*.^79,80^ In addition, there is a debate on the expression levels of *nr3c2* during the early development depending on the quantitative methods.^21,81^ While there are several investigations attesting central and behavioral roles of MR,^49,75,76,82^ GR is deemed to be involved in the phasic response of SR during circadian peaks of cortisol levels and rapid stress response.^3,10,83^ In our investigation, homozygous *nr3c2* mutants did not diverge in locomotor levels from their WT siblings in any of the assays that we performed: 1-minute light assay,^15^ 2- and 4-minute dark-light repeat assays (Fig. 7), and even in the hyperosmotic stress assays.^15^ On the contrary, other groups reported functional roles that MR plays in HPI axis modulation and resultant behavioral outcomes in teleost fish.^49,76,84^ Such variance may arise from differences in husbandry practices. We maintain all our WT and mutant allele-carrying stock via outbreeding with unrelated WT families. Homozygous mutant larvae were spawned only for the experimental purposes by breeding the heterozygous parental pair and euthanized after assays were done. However, in some other investigations, homozygous larvae were obtained from homozygous adult fish (maternal-zygotic). Zebrafish do not tolerate recessive homozygous alleles and there is an increased risk to have aberrant phenotypes after a few generations of inbreeding.^85^

### Investigating adaptive behavior may help discover genetic modifiers of vertebrate stress response

We reported adaptive locomotor response that depended on the length and intensity of light. For the behavior, canonical HPA axis activity (*mc2r, nr3c1*) was required after shorter light exposure but was dispensable after longer illumination. In addition to the light assay paradigms, we previously showed that HPA axis-dependent behavior can be studied, utilizing the osmoregulatory process.^15^ Since HPA axis-mediated vertebrate stress response modulates a wide range of physiological processes, multiple modalities of behavioral assays would enable us to investigate distinct facets of stress response and core pathways redundant in multiple processes (Fig. 9B). For example, feeding states was shown to alter animal behaviors.^86^ Changing levels of environmental light are correlated with mood disorders in humans and behaviors in animals.^87–90^ With the behavioral screening, we hope to discover novel genetic modifiers of the stress response and further our understanding on the core processes of this vertebrate-specific phenomenon.

## Supporting information

Supplemental Result 1

## Acknowledgements

We appreciate the staff of the Mayo Clinic Library for their literature search service and the National Institutes of Health for funding this project (R01 GM134732).

## References

1 Chinenov, Y., Gupte, R. & Rogatsky, I. Nuclear receptors in inflammation control: repression by GR and beyond. Mol Cell Endocrinol 380, 55–64, doi:10.1016/j.mce.2013.04.006 (2013).

2 Selye, H. Stress and the general adaptation syndrome. Br Med J 1, 1383–1392 (1950).

3 Russell, G. & Lightman, S. The human stress response. Nat Rev Endocrinol 15, 525–534, doi:10.1038/s41574-019-0228-0 (2019).

4 Herman, J. P. et al. Regulation of the Hypothalamic-Pituitary-Adrenocortical Stress Response. Compr Physiol 6, 603–621, doi:10.1002/cphy.c150015 (2016).

5 Clark, A. J. L. & Chan, L. Stability and Turnover of the ACTH Receptor Complex. Front Endocrinol (Lausanne) 10, 491, doi:10.3389/fendo.2019.00491 (2019).

6 Arnett, M. G., Muglia, L. M., Laryea, G. & Muglia, L. J. Genetic Approaches to Hypothalamic-Pituitary-Adrenal Axis Regulation. Neuropsychopharmacology 41, 245–260, doi:10.1038/npp.2015.215 (2016).

7 DeMorrow, S. Role of the Hypothalamic-Pituitary-Adrenal Axis in Health and Disease. Int J Mol Sci 19, doi:10.3390/ijms19040986 (2018).

8 Rankin, J., Walker, J. J., Windle, R., Lightman, S. L. & Terry, J. R. Characterizing dynamic interactions between ultradian glucocorticoid rhythmicity and acute stress using the phase response curve. PLoS One 7, e30978, doi:10.1371/journal.pone.0030978 (2012).

9 Sarabdjitsingh, R. A. et al. Stress responsiveness varies over the ultradian glucocorticoid cycle in a brain-region-specific manner. Endocrinology 151, 5369–5379, doi:10.1210/en.2010-0832 (2010).

10 Lightman, S. L. & Conway-Campbell, B. L. The crucial role of pulsatile activity of the HPA axis for continuous dynamic equilibration. Nat Rev Neurosci 11, 710–718, doi:10.1038/nrn2914 (2010).

11 de Kloet, E. R., Meijer, O. C., de Nicola, A. F., de Rijk, R. H. & Joels, M. Importance of the brain corticosteroid receptor balance in metaplasticity, cognitive performance and neuro-inflammation. Front Neuroendocrinol 49, 124–145, doi:10.1016/j.yfrne.2018.02.003 (2018).

12 de Kloet, E. R. From receptor balance to rational glucocorticoid therapy. Endocrinology 155, 2754–2769, doi:10.1210/en.2014-1048 (2014).

13 Joels, M., Pasricha, N. & Karst, H. The interplay between rapid and slow corticosteroid actions in brain. Eur J Pharmacol 719, 44–52, doi:10.1016/j.ejphar.2013.07.015 (2013).

14 De Kloet, E. R., Vreugdenhil, E., Oitzl, M. S. & Joels, M. Brain corticosteroid receptor balance in health and disease. Endocr Rev 19, 269–301, doi:10.1210/edrv.19.3.0331 (1998).

15 Lee, H. B. et al. Novel zebrafish behavioral assay to identify modifiers of the rapid, nongenomic stress response. Genes Brain Behav 18, e12549, doi:10.1111/gbb.12549 (2019).

16 Schaaf, M. J., Chatzopoulou, A. & Spaink, H. P. The zebrafish as a model system for glucocorticoid receptor research. Comp Biochem Physiol A Mol Integr Physiol 153, 75–82, doi:10.1016/j.cbpa.2008.12.014 (2009).

17 Facchinello, N. et al. nr3c1 null mutant zebrafish are viable and reveal DNA-binding-independent activities of the glucocorticoid receptor. Sci Rep 7, 4371, doi:10.1038/s41598-017-04535-6 (2017).

18 Spence, R., Gerlach, G., Lawrence, C. & Smith, C. The behaviour and ecology of the zebrafish, Danio rerio. Biol Rev Camb Philos Soc 83, 13–34, doi:10.1111/j.1469-185X.2007.00030.x (2008).

19 Kimmel, C. B., Ballard, W. W., Kimmel, S. R., Ullmann, B. & Schilling, T. F. Stages of embryonic development of the zebrafish. Dev Dyn 203, 253–310, doi:10.1002/aja.1002030302 (1995).

20 Alsop, D. & Vijayan, M. The zebrafish stress axis: molecular fallout from the teleost-specific genome duplication event. Gen Comp Endocrinol 161, 62–66, doi:10.1016/j.ygcen.2008.09.011 (2009).

21 Alsop, D. & Vijayan, M. M. Development of the corticosteroid stress axis and receptor expression in zebrafish. Am J Physiol Regul Integr Comp Physiol 294, R711–719, doi:10.1152/ajpregu.00671.2007 (2008).

22 Nesan, D. & Vijayan, M. M. Role of glucocorticoid in developmental programming: evidence from zebrafish. Gen Comp Endocrinol 181, 35–44, doi:10.1016/j.ygcen.2012.10.006 (2013).

23 Fernandes, A. M. et al. Deep brain photoreceptors control light-seeking behavior in zebrafish larvae. Curr Biol 22, 2042–2047, doi:10.1016/j.cub.2012.08.016 (2012).

24 Burgess, H. A., Schoch, H. & Granato, M. Distinct retinal pathways drive spatial orientation behaviors in zebrafish navigation. Curr Biol 20, 381–386, doi:10.1016/j.cub.2010.01.022 (2010).

25 Horstick, E. J., Bayleyen, Y., Sinclair, J. L. & Burgess, H. A. Search strategy is regulated by somatostatin signaling and deep brain photoreceptors in zebrafish. BMC Biol 15, 4, doi:10.1186/s12915-016-0346-2 (2017).

26 Brockerhoff, S. E. et al. A behavioral screen for isolating zebrafish mutants with visual system defects. Proc Natl Acad Sci U S A 92, 10545–10549, doi:10.1073/pnas.92.23.10545 (1995).

27 Burgess, H. A. & Granato, M. Modulation of locomotor activity in larval zebrafish during light adaptation. J Exp Biol 210, 2526–2539, doi:10.1242/jeb.003939 (2007).

28 Paul, K. N., Saafir, T. B. & Tosini, G. The role of retinal photoreceptors in the regulation of circadian rhythms. Rev Endocr Metab Disord 10, 271–278, doi:10.1007/s11154-009-9120-x (2009).

29 Oster, H. The interplay between stress, circadian clocks, and energy metabolism. Journal of Endocrinology 247, R13–R25, doi:10.1530/joe-20-0124 (2020).

30 Minnetti, M. et al. Fixing the broken clock in adrenal disorders: focus on glucocorticoids and chronotherapy. J Endocrinol 246, R13–R31, doi:10.1530/JOE-20-0066 (2020).

31 Jaikumar, G., Slabbekoorn, H., Sireeni, J., Schaaf, M. & Tudorache, C. The role of the Glucocorticoid Receptor in the Regulation of Diel Rhythmicity. Physiol Behav 223, 112991, doi:10.1016/j.physbeh.2020.112991 (2020).

32 Morbiato, E. et al. Feeding Entrainment of the Zebrafish Circadian Clock Is Regulated by the Glucocorticoid Receptor. Cells 8, doi:10.3390/cells8111342 (2019).

33 Ruginsk, S. G. et al. in Corticosteroids Ch. Chapter 3, (2018).

34 Muto, A. et al. Forward genetic analysis of visual behavior in zebrafish. PLoS Genet 1, e66, doi:10.1371/journal.pgen.0010066 (2005).

35 Muto, A., Taylor, M. R., Suzawa, M., Korenbrot, J. I. & Baier, H. Glucocorticoid receptor activity regulates light adaptation in the zebrafish retina. Front Neural Circuits 7, 145, doi:10.3389/fncir.2013.00145 (2013).

36 Nüsslein-Volhard, C. & Dahm, R. Zebrafish: A practical approach. 1st edn, (Oxford University Press, 2002).

37 Uchida, D., Yamashita, M., Kitano, T. & Iguchi, T. Oocyte apoptosis during the transition from ovary-like tissue to testes during sex differentiation of juvenile zebrafish. J Exp Biol 205, 711–718, doi:10.1242/jeb.205.6.711 (2002).

38 Takahashi, H. Juvenile hermaphroditism in the zebrafish, Brachydanio rerio. Bull Fac Fish Hokkaido Univ 28, 57–65 (1977).

39 Orban, L., Sreenivasan, R. & Olsson, P. E. Long and winding roads: testis differentiation in zebrafish. Mol Cell Endocrinol 312, 35–41, doi:10.1016/j.mce.2009.04.014 (2009).

40 Hartmann, S. et al. Zebrafish larvae show negative phototaxis to near-infrared light. PLoS One 13, e0207264, doi:10.1371/journal.pone.0207264 (2018).

41 R: A language and environment for statistical computing. R Foundation for Statistical Computing (2021).

42 MacPhail, R. C. et al. Locomotion in larval zebrafish: Influence of time of day, lighting and ethanol. Neurotoxicology 30, 52–58, doi:10.1016/j.neuro.2008.09.011 (2009).

43 Ellis, L. D., Seibert, J. & Soanes, K. H. Distinct models of induced hyperactivity in zebrafish larvae. Brain Res 1449, 46–59, doi:10.1016/j.brainres.2012.02.022 (2012).

44 Kokel, D. et al. Identification of nonvisual photomotor response cells in the vertebrate hindbrain. J Neurosci 33, 3834–3843, doi:10.1523/JNEUROSCI.3689-12.2013 (2013).

45 Kokel, D. et al. Rapid behavior-based identification of neuroactive small molecules in the zebrafish. Nat Chem Biol 6, 231–237, doi:10.1038/nchembio.307 (2010).

46 Haigis, A. C., Ottermanns, R., Schiwy, A., Hollert, H. & Legradi, J. Getting more out of the zebrafish light dark transition test. Chemosphere 295, 133863, doi:10.1016/j.chemosphere.2022.133863 (2022).

47 Colon-Cruz, L. et al. Alterations of larval photo-dependent swimming responses (PDR): New endpoints for rapid and diagnostic screening of aquatic contamination. Ecotoxicol Environ Saf 147, 670–680, doi:10.1016/j.ecoenv.2017.09.018 (2018).

48 Irons, T. D., Kelly, P. E., Hunter, D. L., Macphail, R. C. & Padilla, S. Acute administration of dopaminergic drugs has differential effects on locomotion in larval zebrafish. Pharmacol Biochem Behav 103, 792–813, doi:10.1016/j.pbb.2012.12.010 (2013).

49 Faught, E. & Vijayan, M. M. The mineralocorticoid receptor is essential for stress axis regulation in zebrafish larvae. Sci Rep 8, 18081, doi:10.1038/s41598-018-36681-w (2018).

50 Bitran, D., Shiekh, M., Dowd, J. A., Dugan, M. M. & Renda, P. Corticosterone is permissive to the anxiolytic effect that results from the blockade of hippocampal mineralocorticoid receptors. Pharmacol Biochem Behav 60, 879–887, doi:10.1016/s0091-3057(98)00071-9 (1998).

51 Anagnostaras, S. G. et al. Automated assessment of pavlovian conditioned freezing and shock reactivity in mice using the video freeze system. Front Behav Neurosci 4, doi:10.3389/fnbeh.2010.00158 (2010).

52 Egan, R. J. et al. Understanding behavioral and physiological phenotypes of stress and anxiety in zebrafish. Behav Brain Res 205, 38–44, doi:10.1016/j.bbr.2009.06.022 (2009).

53 Kalueff, A. V., Kaluyeva, A. & Maillet, E. L. Anxiolytic-like effects of noribogaine in zebrafish. Behav Brain Res 330, 63–67, doi:10.1016/j.bbr.2017.05.008 (2017).

54 Best, C. & Vijayan, M. M. Cortisol elevation post-hatch affects behavioural performance in zebrafish larvae. Gen Comp Endocrinol 257, 220–226, doi:10.1016/j.ygcen.2017.07.009 (2018).

55 Best, C., Kurrasch, D. M. & Vijayan, M. M. Maternal cortisol stimulates neurogenesis and affects larval behaviour in zebrafish. Sci Rep 7, 40905, doi:10.1038/srep40905 (2017).

56 Golla, A., Ostby, H. & Kermen, F. Chronic unpredictable stress induces anxiety-like behaviors in young zebrafish. Sci Rep 10, 10339, doi:10.1038/s41598-020-67182-4 (2020).

57 Packard, A. E., Egan, A. E. & Ulrich-Lai, Y. M. HPA Axis Interactions with Behavioral Systems. Compr Physiol 6, 1897–1934, doi:10.1002/cphy.c150042 (2016).

58 Deroche, V. et al. Stress-induced sensitization to amphetamine and morphine psychomotor effects depend on stress-induced corticosterone secretion. Brain Res 598, 343–348, doi:10.1016/0006-8993(92)90205-n (1992).

59 Roberts, A. J., Lessov, C. N. & Phillips, T. J. Critical role for glucocorticoid receptors in stress- and ethanol-induced locomotor sensitization. J Pharmacol Exp Ther 275, 790–797 (1995).

60 Schumacher, M., Coirini, H., Pfaff, D. W. & McEwen, B. S. Behavioral effects of progesterone associated with rapid modulation of oxytocin receptors. Science 250, 691–694, doi:10.1126/science.2173139 (1990).

61 Schumacher, M. Rapid membrane effects of steroid hormones: an emerging concept in neuroendocrinology. Trends Neurosci 13, 359–362, doi:10.1016/0166-2236(90)90016-4 (1990).

62 Smythe, J. W., Murphy, D., Timothy, C. & Costall, B. Hippocampal mineralocorticoid, but not glucocorticoid, receptors modulate anxiety-like behavior in rats. Pharmacol Biochem Behav 56, 507–513, doi:10.1016/s0091-3057(96)00244-4 (1997).

63 De Kloet, E. R. Hormones and the stressed brain. Ann N Y Acad Sci 1018, 1–15, doi:10.1196/annals.1296.001 (2004).

64 Hartig, E. I., Zhu, S., King, B. L. & Coffman, J. A. Chronic cortisol exposure in early development leads to neuroendocrine dysregulation in adulthood. BMC Res Notes 13, 366, doi:10.1186/s13104-020-05208-w (2020).

65 Laryea, G., Muglia, L., Arnett, M. & Muglia, L. J. Dissection of glucocorticoid receptor-mediated inhibition of the hypothalamic-pituitary-adrenal axis by gene targeting in mice. Front Neuroendocrinol 36, 150–164, doi:10.1016/j.yfrne.2014.09.002 (2015).

66 Duma, D., Jewell, C. M. & Cidlowski, J. A. Multiple glucocorticoid receptor isoforms and mechanisms of post-translational modification. J Steroid Biochem Mol Biol 102, 11–21, doi:10.1016/j.jsbmb.2006.09.009 (2006).

67 Lu, N. Z. & Cidlowski, J. A. Glucocorticoid receptor isoforms generate transcription specificity. Trends Cell Biol 16, 301–307, doi:10.1016/j.tcb.2006.04.005 (2006).

68 Oakley, R. H. & Cidlowski, J. A. Cellular processing of the glucocorticoid receptor gene and protein: new mechanisms for generating tissue-specific actions of glucocorticoids. J Biol Chem 286, 3177–3184, doi:10.1074/jbc.R110.179325 (2011).

69 Chida, D. et al. Melanocortin 2 receptor is required for adrenal gland development, steroidogenesis, and neonatal gluconeogenesis. Proc Natl Acad Sci U S A 104, 18205–18210, doi:10.1073/pnas.0706953104 (2007).

70 Heym, C. Immunocytochemical correlates of an extrapituitary adrenocortical regulation in man. Histol Histopathol 12, 567–581, doi:10.14670/HH-12.567 (1997).

71 Hsu, H. J., Lin, G. & Chung, B. C. Parallel early development of zebrafish interrenal glands and pronephros: differential control by wt1 and ff1b. Development 130, 2107–2116, doi:10.1242/dev.00427 (2003).

72 Liu, Y. W. Interrenal organogenesis in the zebrafish model. Organogenesis 3, 44–48, doi:10.4161/org.3.1.3965 (2007).

73 Josep Agulleiro, M. et al. Melanocortin 4 receptor becomes an ACTH receptor by coexpression of melanocortin receptor accessory protein 2. Mol Endocrinol 27, 1934–1945, doi:10.1210/me.2013-1099 (2013).

74 Alderman, S. L. & Vijayan, M. M. 11beta-Hydroxysteroid dehydrogenase type 2 in zebrafish brain: a functional role in hypothalamus-pituitary-interrenal axis regulation. J Endocrinol 215, 393–402, doi:10.1530/JOE-12-0379 (2012).

75 Sturm, A. et al. 11-deoxycorticosterone is a potent agonist of the rainbow trout (Oncorhynchus mykiss) mineralocorticoid receptor. Endocrinology 146, 47–55, doi:10.1210/en.2004-0128 (2005).

76 Takahashi, H. & Sakamoto, T. The role of ‘mineralocorticoids’ in teleost fish: relative importance of glucocorticoid signaling in the osmoregulation and ‘central’ actions of mineralocorticoid receptor. Gen Comp Endocrinol 181, 223–228, doi:10.1016/j.ygcen.2012.11.016 (2013).

77 Tokarz, J., Norton, W., Moller, G., Hrabe de Angelis, M. & Adamski, J. Zebrafish 20beta-hydroxysteroid dehydrogenase type 2 is important for glucocorticoid catabolism in stress response. PLoS One 8, e54851, doi:10.1371/journal.pone.0054851 (2013).

78 McCormick, S. D., Regish, A., O’Dea, M. F. & Shrimpton, J. M. Are we missing a mineralocorticoid in teleost fish? Effects of cortisol, deoxycorticosterone and aldosterone on osmoregulation, gill Na+,K+ -ATPase activity and isoform mRNA levels in Atlantic salmon. Gen Comp Endocrinol 157, 35–40, doi:10.1016/j.ygcen.2008.03.024 (2008).

79 Cruz, S. A., Lin, C. H., Chao, P. L. & Hwang, P. P. Glucocorticoid receptor, but not mineralocorticoid receptor, mediates cortisol regulation of epidermal ionocyte development and ion transport in zebrafish (danio rerio). PLoS One 8, e77997, doi:10.1371/journal.pone.0077997 (2013).

80 Kumai, Y., Nesan, D., Vijayan, M. M. & Perry, S. F. Cortisol regulates Na+ uptake in zebrafish, Danio rerio, larvae via the glucocorticoid receptor. Mol Cell Endocrinol 364, 113–125, doi:10.1016/j.mce.2012.08.017 (2012).

81 Bertrand, S. et al. Unexpected novel relational links uncovered by extensive developmental profiling of nuclear receptor expression. PLoS Genet 3, e188, doi:10.1371/journal.pgen.0030188 (2007).

82 Trayer, V., Hwang, P. P., Prunet, P. & Thermes, V. Assessment of the role of cortisol and corticosteroid receptors in epidermal ionocyte development in the medaka (Oryzias latipes) embryos. Gen Comp Endocrinol 194, 152–161, doi:10.1016/j.ygcen.2013.09.011 (2013).

83 Gjerstad, J. K., Lightman, S. L. & Spiga, F. Role of glucocorticoid negative feedback in the regulation of HPA axis pulsatility. Stress 21, 403–416, doi:10.1080/10253890.2018.1470238 (2018).

84 Sakamoto, T. et al. Principal function of mineralocorticoid signaling suggested by constitutive knockout of the mineralocorticoid receptor in medaka fish. Sci Rep 6, 37991, doi:10.1038/srep37991 (2016).

85 Lee, H. B., Modhurima, R., Heeren, A. A. & Clark, K. J. in Behavioral and Neural Genetics of Zebrafish 263–278 (2020).

86 Filosa, A., Barker, A. J., Dal Maschio, M. & Baier, H. Feeding State Modulates Behavioral Choice and Processing of Prey Stimuli in the Zebrafish Tectum. Neuron 90, 596–608, doi:10.1016/j.neuron.2016.03.014 (2016).

87 Juruena, M. F. & Cleare, A. J. Overlap between atypical depression, seasonal affective disorder and chronic fatigue syndrome. Braz J Psychiatry 29 Suppl 1, S19–26, doi:10.1590/s1516-44462007000500005 (2007).

88 Lonstein, J. S., Linning-Duffy, K. & Yan, L. Low Daytime Light Intensity Disrupts Male Copulatory Behavior, and Upregulates Medial Preoptic Area Steroid Hormone and Dopamine Receptor Expression, in a Diurnal Rodent Model of Seasonal Affective Disorder. Front Behav Neurosci 13, 72, doi:10.3389/fnbeh.2019.00072 (2019).

89 Bao, A. M. & Swaab, D. F. The human hypothalamus in mood disorders: The HPA axis in the center. IBRO Rep 6, 45–53, doi:10.1016/j.ibror.2018.11.008 (2019).

90 Ziv, L. et al. An affective disorder in zebrafish with mutation of the glucocorticoid receptor. Mol Psychiatry 18, 681–691, doi:10.1038/mp.2012.64 (2013).

