## Supplemental Result 1 for "The canonical HPA axis contributes to locomotion during photoadaptation but is not required"

**Supplemental Result 1. Statistical analyses**

**Figure 2. Loss of *nr3c1* larvae exhibited decreased basal locomotor activity in lit and dark environments.**

(A) WT baseline assay: A three-way mixed ANOVA was performed with ambient light (darkness for 12 hrs vs. lit environment for 12 hrs) as the between-subject independent variable, larval age (4, 5, 6, and 7 dpf) as the between-subject independent variable, and time of the day (10 am – 12 pm, 12 – 2 pm, 2 – 4 pm, 4 – 6 pm, 6 – 8 pm, and 8 – 10 pm) as the within-subject independent variable.

There was a significant main effect of ambient light on locomotor levels, $F(1, 2104) = 762.97$, $p < 0.0001$. There was a significant main effect of larval age, $F(3, 2104) = 257.38$, $p < 0.0001$. There was a significant main effect of time of the day, $F(5, 2100) = 108.23$, $p < 0.0001$.

There was a significant ambient light x larval age interaction on locomotor levels, $F(3, 2104) = 98.14$, $p < 0.0001$. There was a significant ambient light x time of the day interaction, $F(5, 2100) = 43.74$, $p < 0.0001$. There was a significant larval age x time of the day interaction, $F(15, 6306) = 19.48$, $p < 0.0001$.

There was a significant ambient light x larval age x time of the day interaction, $F(15, 6306) = 12.96$, $p < 0.0001$. The locomotor levels differed based on the ambient light condition in which larvae were video recorded, based on the larval age and the time of the day.

The degrees of freedom of the error term were adjusted after Greenhouse-Geisser correction.

(B) nr3c1 exon 5 baseline assay: A three-way mixed ANOVA was performed with ambient light (darkness for 12 hrs vs. lit environment for 12 hrs) as the between-subject independent variable, genotype (WT, het, and hom) as the between-subject independent variable, and time of the day (10 am – 12 pm, 12 – 2 pm, 2 – 4 pm, 4 – 6 pm, 6 – 8 pm, and 8 – 10 pm) as the within-subject independent variable.

There was a significant main effect of ambient light on locomotor levels, $F(1, 560) = 47.44$, $p < 0.0001$. There was a significant main effect of genotype, $F(2, 560) = 8.93$, $p = 0.0002$. There was a significant main effect of time of the day, $F(5, 556) = 31.41$, $p < 0.0001$.

There was a significant ambient light x genotype interaction on locomotor levels, $F(2, 560) = 3.38$, $p = 0.035$. There was a significant ambient light x time of the day interaction, $F(5, 556) = 4.12$, $p = 0.001$. There was a significant genotype x time of the day interaction, $F(10, 1114) = 2.40$, $p < 0.008$.

The interaction among ambient light x larval age x time of the day was not significant, $F(10, 1114) = 1.39$, $p = 0.18$. The locomotor levels differed based on the ambient light in which larvae were video recorded and the time of the day. The homozygous mutants showed decreased locomotion in several segments in the time of the day.

The degrees of freedom of the error term were adjusted after Greenhouse-Geisser correction.

**Figure 3. Mutant *mc2r* and *nr3c1* larvae showed locomotion indistinguishable from their WT siblings in 7.5-min repeat assays.**

(A) *mc2r* 7.5-min repeat: A two-way mixed ANOVA was performed with genotype (WT, het, and hom) as the between-subject independent variable and illumination (dark, light, dark, light, dark, light, dark, light) as the within-subject independent variable.

There was a significant main effect of genotype on locomotor levels, $F(2, 548) = 8.44$, $p = 0.0002$. There was a significant main effect of illumination, $F(7, 542) = 82.53$, $p < 0.0001$. There was a significant genotype x illumination interaction on locomotor levels, $F(14, 1086) = 3.38$, $p < 0.0001$. *mc2r* homozygous mutants showed significantly decreased locomotion during some of the dark phase. The degrees of freedom of the error term were adjusted after Greenhouse-Geisser correction.

(B) *nr3c1* exon 2 7.5-min repeat: A two-way mixed ANOVA was performed with genotype (WT, het, and hom) as the between-subject independent variable and illumination (dark, light, dark, light, dark, light, dark, light) as the within-subject independent variable.

The main effect of genotype on locomotor levels was not significant, $F(2, 544) = 1.85$, $p = 0.16$. There was a significant main effect of illumination, $F(7, 538) = 187.27$, $p < 0.0001$. No significant genotype x illumination interaction on locomotor levels was found, $F(14, 1078) = 0.74$, $p = 0.74$. The degrees of freedom of the error term were adjusted after Greenhouse-Geisser correction.

(C) *nr3c1* exon 5 7.5-min repeat: A two-way mixed ANOVA was performed with genotype (WT, het, and hom) as the between-subject independent variable and illumination (dark, light, dark, light, dark, light, dark, light) as the within-subject independent variable.

The main effect of genotype on locomotor levels was not significant, $F(2, 1161) = 1.79$, $p = 0.17$. There was a significant main effect of illumination, $F(7, 1155) = 85.83$, $p < 0.0001$. No significant genotype x illumination interaction on locomotor levels was found, $F(14, 2312) = 0.90$, $p = 0.56$. The degrees of freedom of the error term were adjusted after Greenhouse-Geisser correction.

**Figure 4. Mutant *mc2r* larvae exhibited decreased locomotion in 4- or 2-min light illumination.**

(A) *mc2r* 7.5-4 min repeat: A two-way mixed ANOVA was performed with genotype (WT, het, and hom) as the between-subject independent variable and illumination (dark, light, dark, light, dark, light, dark, light, dark) as the within-subject independent variable.

There was a significant main effect of genotype on locomotor levels, $F(2, 580) = 18.60$, $p < 0.0001$. There was a significant main effect of illumination, $F(8, 573) = 80.96$, $p < 0.0001$. There was a significant genotype x illumination interaction on locomotor levels, $F(16, 1148) = 3.30$, $p < 0.0001$. *mc2r* homozygous mutants showed significantly decreased locomotion in all of the dark phase. The degrees of freedom of the error term were adjusted after Greenhouse-Geisser correction.

(B) *mc2r* 7.5-2 min repeat: A two-way mixed ANOVA was performed with genotype (WT, het, and hom) as the between-subject independent variable and illumination (dark, light, dark, light, dark, light, dark, light, dark) as the within-subject independent variable.

There was a significant main effect of genotype on locomotor levels, $F(2, 947) = 28.87$, $p < 0.0001$. There was a significant main effect of illumination, $F(8, 940) = 100.57$, $p < 0.0001$. There was a significant genotype x illumination interaction on locomotor levels, $F(16, 1882) = 5.96$, $p < 0.0001$. *mc2r* homozygous mutants showed significantly decreased locomotion in all of the dark phase. The degrees of freedom of the error term were adjusted after Greenhouse-Geisser correction.

**Figure 5. Mutant *nr3c1* (exon 2) larvae showed decreasing trends in locomotion when subjected to 6-, or 4-min of light.**

(A) *nr3c1* exon 2 7.5-6 min repeat: A two-way mixed ANOVA was performed with genotype (WT, het, and hom) as the between-subject independent variable and illumination (dark, light, dark, light, dark, light, dark, light, dark) as the within-subject independent variable.

The main effect of genotype on locomotor levels was not significant, $F(2, 271) = 1.95$, $p = 0.14$. There was a significant main effect of illumination, $F(8, 264) = 74.80$, $p < 0.0001$. There was a significant genotype x illumination interaction on locomotor levels, $F(16, 530) = 2.08$, $p = 0.008$. The degrees of freedom of the error term were adjusted after Greenhouse-Geisser correction.

(B) *nr3c1* exon 2 7.5-4 min repeat: A two-way mixed ANOVA was performed with genotype (WT, het, and hom) as the between-subject independent variable and illumination (dark, light, dark, light, dark, light, dark, light, dark) as the within-subject independent variable.

There was a significant main effect of genotype on locomotor levels, $F(2, 651) = 8.97$, $p = 0.0001$. There was a significant main effect of illumination, $F(8, 644) = 110.57$, $p < 0.0001$. There was a significant genotype x illumination interaction on locomotor levels, $F(16, 1290) = 2.17$, $p = 0.005$. *nr3c1* homozygous mutants showed significantly decreased locomotion in some of the dark phase. The degrees of freedom of the error term were adjusted after Greenhouse-Geisser correction.

**Figure 6. Mutant *nr3c1* (exon 5) larvae displayed decreasing trends in locomotion when subjected to 6-, 4-, or 2-min of light.**

(A) *nr3c1* exon 5 7.5-6 min repeat: A two-way mixed ANOVA was performed with genotype (WT, het, and hom) as the between-subject independent variable and illumination (dark, light, dark, light, dark, light, dark, light, dark) as the within-subject independent variable.

There was a significant main effect of genotype on locomotor levels, $F(2, 559) = 9.31$, $p = 0.0001$. There was a significant main effect of illumination, $F(8, 552) = 79.94$, $p < 0.0001$. There was a significant genotype x illumination interaction on locomotor levels, $F(16, 1106) = 1.85$, $p = 0.022$. *nr3c1* homozygous mutants showed significantly decreased locomotion in some of the dark phase. The degrees of freedom of the error term were adjusted after Greenhouse-Geisser correction.

(B) *nr3c1* exon 5 7.5-4 min repeat: A two-way mixed ANOVA was performed with genotype (WT, het, and hom) as the between-subject independent variable and illumination (dark, light, dark, light, dark, light, dark, light, dark) as the within-subject independent variable.

There was a significant main effect of genotype on locomotor levels, $F(2, 311) = 7.64$, $p = 0.0006$. There was a significant main effect of illumination, $F(8, 304) = 76.27$, $p < 0.0001$. There was a significant genotype x illumination interaction on locomotor levels, $F(16, 610) = 2.71$, $p = 0.0003$. *nr3c1* homozygous mutants showed significantly decreased locomotion in some of the dark phase. The degrees of freedom of the error term were adjusted after Greenhouse-Geisser correction.

(C) *nr3c1* exon 5 7.5-2 min repeat: A two-way mixed ANOVA was performed with genotype (WT, het, and hom) as the between-subject independent variable and illumination (dark, light, dark, light, dark, light, dark, light, dark) as the within-subject independent variable.

There was a significant main effect of genotype on locomotor levels, $F(2, 368) = 21.27$, $p < 0.0001$. There was a significant main effect of illumination, $F(8, 361) = 48.02$, $p < 0.0001$. There was a significant genotype x illumination interaction on locomotor levels, $F(16, 724) = 3.30$, $p < 0.0001$. *nr3c1* homozygous mutants showed significantly decreased locomotion in some of the dark phase. The degrees of freedom of the error term were adjusted after Greenhouse-Geisser correction.

**Figure 7. Mutant *nr3c2* larvae showed locomotion similar to their WT siblings in 7.5-, 4- or 2-min illumination.**

(A) *nr3c2* exon 2 7.5-min repeat: A two-way mixed ANOVA was performed with genotype (WT, het, and hom) as the between-subject independent variable and illumination (dark, light, dark, light, dark, light) as the within-subject independent variable.

The main effect of genotype on locomotor levels was not significant, $F(2, 441) = 0.04$, $p = 0.96$. There was a significant main effect of illumination, $F(7, 435) = 105.52$, $p < 0.0001$. No significant genotype x illumination interaction on locomotor levels was found, $F(14, 872) = 0.70$, $p = 0.77$. The degrees of freedom of the error term were adjusted after Greenhouse-Geisser correction.

(B) *nr3c2* exon 2 7.5-4 min repeat: A two-way mixed ANOVA was performed with genotype (WT, het, and hom) as the between-subject independent variable and illumination (dark, light, dark, light, dark, light, dark) as the within-subject independent variable.

The main effect of genotype on locomotor levels was not significant, $F(2, 267) = 1.18$, $p = 0.31$. There was a significant main effect of illumination, $F(8, 260) = 55.11$, $p < 0.0001$. No significant genotype x illumination interaction on locomotor levels was found, $F(16, 522) = 1.42$, $p = 0.13$. The degrees of freedom of the error term were adjusted after Greenhouse-Geisser correction.

(C) *nr3c2* exon 2 7.5-2 min repeat: A two-way mixed ANOVA was performed with genotype (WT, het, and hom) as the between-subject independent variable and illumination (dark, light, dark, light, dark, light) as the within-subject independent variable.

The main effect of genotype on locomotor levels was not significant, $F(2, 433) = 2.06$, $p = 0.13$. There was a significant main effect of illumination, $F(7, 427) = 103.70$, $p < 0.0001$. No significant genotype x illumination interaction on locomotor levels was found, $F(14, 856) = 1.44$, $p = 0.13$. The degrees of freedom of the error term were adjusted after Greenhouse-Geisser correction.

**Figure 8. Dimmer white light produced similar locomotion in *nr3c1* WT, heterozygous, and homozygous siblings.**

(A) *nr3c1* exon 2 1-min assay: A two-way mixed ANOVA was performed with genotype (WT, het, and hom) as the between-subject independent variable and illumination (dark pre-treatment, dark post-treatment) as the within-subject independent variable.

The main effect of genotype on locomotor levels was not significant, $F(2, 543) = 0.74$, $p = 0.48$. There was a significant main effect of illumination, $F(1, 543) = 19.21$, $p < 0.0001$. No significant genotype x illumination interaction on locomotor levels was found, $F(2, 543) = 0.004$, $p = 1.00$. The degrees of freedom of the error term were adjusted after Greenhouse-Geisser correction.

(B) *nr3c1* exon 5 7.5-7.5 min repeat: A two-way mixed ANOVA was performed with genotype (WT, het, and hom) as the between-subject independent variable and illumination (dark, light, dark, light, dark, light, dark) as the within-subject independent variable.

The main effect of genotype on locomotor levels was not significant, $F(2, 654) = 0.01$, $p = 0.99$. There was a significant main effect of illumination, $F(8, 647) = 154.43$, $p < 0.0001$. No significant genotype x illumination interaction on locomotor levels was found, $F(16, 1296) = 1.41$, $p = 0.13$. The degrees of freedom of the error term were adjusted after Greenhouse-Geisser correction.
