## Supplemental Table 1 for "The canonical HPA axis contributes to locomotion during photoadaptation but is not required"

### Supplemental Table 1. Materials and equipment

| Materials/Instruments | Cat. No | Vendor | Address |
|---|---|---|---|
| 2-amino-2-(hydroxymethyl)-1,3-propanediol (Tris) | T1378 | Sigma-Aldrich Co. LLC. | St. Louis, MO, USA |
| 48-well plates | 08-772-1C | Falcon | Waltham, MA, USA |
| 5x MyTaq™ Reaction Buffer Colorless | BIO-37111 | Bioline USA Inc. | Taunton, ME, USA |
| 5x MyTaqTM Red Reaction Buffer | BIO-37112 | Bioline USA Inc. | Taunton, ME, USA |
| ASQ Fluorescent probes and quenchers | - | IDT, Inc. | Coralville, IA, USA |
| Baffle / Screen (400 µm) | ZT280S400 | Aquaneering, Inc. | San Diego, CA, USA |
| Benchtop Optical Power Meter | 1936-R | Newport Corp. | Irvine, CA, USA |
| Breeding tanks (1 L) | ZB17BTE/L/D/IS LOP | Tecniplast, Inc. | West Chester, PA, USA |
| CFX96 C1000 Touch™ | 185-5096 | Bio-Rad Laboratories, Inc. | Hercules, CA, USA |
| Cortisol ELISA Kit | 500360 | Cayman Chemical | Ann Arbor, MI, USA |
| DC12V SMD3528-600-IR InfraRed Single Chip Flexible LED Strips 120L EDs 9.6W Per Meter | HK-F3528IR60-X | LED Lights World LTD. | Shenzhen, GuangDong, China |
| Diethyl ether | AC615080010 | Acros organics | Waltham, MA, USA |
| Ethylenediaminetetraacetic acid disodium salt dihydrate (EDTA) | E-5134 | Sigma-Aldrich Co. LLC. | St. Louis, MO, USA |
| Handycam Camera | HDR-CX560V | Sony Corp. | New York City, NY, USA |
| Housing tanks (3, 9 L) | ZT280, 950 | Aquaneering, Inc. | San Diego, CA, USA |
| Light box with white LED strips (40W)* | LPW-xW6060-40 | Super Bright LEDs Inc. | St. Louis, MO, USA |
| Light boxes* | - | Division of Engineering, Mayo Clinic | Rochester, MN, USA |
| Pico liter injector (PLI-90) | EC1 65-0004 | Harvard Apparatus | Holliston, MA, USA |
| Microseal® 'B' seal | MSB1001 | Bio-Rad Laboratories, Inc. | Hercules, CA, USA |
| Miniature spectrometer | STS-VIS & STS-NIR | Ocean Optics Inc. | Largo, FL, USA |
| mMESSAGE mMACHINE® T3 Transcription Kit | AM1348 | Ambion | Waltham, MA, USA |
| Multiplate™ PCR plates 96-well, clear | MLL9601 | Bio-Rad Laboratories, Inc. | Hercules, CA, USA |
| MyTaq™ DNA Polymerase | BIO-21106 | Bioline USA Inc. | Taunton, ME, USA |
| MyTaq™ HS DNA Polymerase (5 units/µL) | BIO-21111 | Bioline USA Inc. | Taunton, ME, USA |
| NanoDrop 2000 | - | Thermo Fisher Scientific Inc. | Waltham, MA, USA |
| Pierce™ BCA Protein Assay Kit | 23225 | Thermo Fisher Scientific, Inc. | Waltham, MA, USA |
| Random primers | 48190011 | Thermo Fisher Scientific, Inc. | Waltham, MA, USA |
| Restriction endonuclease | - | New England Biolabs, Inc. | Ipswich, MA, USA |
| Round gel loading tip | NC9531852 | Fisher scientific | Waltham, MA, USA |
| SensiFAST™ SYBR® No-ROX Kit | BIO-98005 | Bioline USA Inc. | Taunton, ME, USA |
| Microloader pipette tips | E5242956003 | Fisher Scientific | Waltham, MA, USA |
| Sodium hydroxide, Pellets (NaOH) | 7708 | Mallinckrodt Limited | Mulhuddart, Dublin 15, Ireland |
| SuperScript™ II Reverse Transcriptase | 18064014 | Invitrogen | Waltham, MA, USA |
| T100™ Thermal Cycler | 186-1096 | Bio-Rad Laboratories, Inc. | Hercules, CA, USA |
| TaqMan Probes™ | - | Applied Biosystems™ | Waltham, MA, USA |
| Vibra-Cell VCX 130 (sonicator) | VCX 130 | Sonics and Materials Inc. | Newtown, CT, USA |
| VorTemp™ 56 shaking incubator | S2056A | Labnet International, Inc. | Edison, NJ, USA |
